# A metallo-beta-lactamase with both beta-lactamase and ribonuclease activity is linked with traduction in giant viruses

**DOI:** 10.1101/819797

**Authors:** Philippe Colson, Lucile Pinault, Said Azza, Nicholas Armstrong, Eric Chabriere, Bernard La Scola, Pierre Pontarotti, Didier Raoult

## Abstract

Enzymatic proteins with a metallo-beta-lactamase (MBL) fold have been essentially studied in bacteria for their activity on beta-lactam antibiotics. However, the MBL fold is ancient and highly conserved, and these proteins are capable of cleaving a broad range of substrates. It has recently been shown that MBLs are present in a wide array of cellular organisms, including eukaryotes and archaea. We show here that Tupanvirus deep ocean, a giant virus, also encodes a protein with a MBL fold. Phylogeny showed its clustering with transfer ribonucleases (RNases) and the presence of orthologs in other giant viruses, mainly those harboring the largest sets of translation components. In addition, it suggests an ancient origin for these genes and a transfer between giant viruses and *Acanthamoeba* spp., a host of many giant viruses. Biologically, after its expression in *Escherichia coli*, the tupanvirus protein was found to hydrolyse nitrocefin, a chromogenic beta-lactam. We also observed an hydrolysis of penicillin G (10 μg/mL) and detected the metabolite of penicillin G hydrolysis, benzylpenilloic acid. This was inhibited by sulbactam, a beta-lactamase inhibitor. In addition, we tested the degradation of single-stranded DNA, double-stranded DNA, and RNAs, and observed a strong activity on RNAs from seven bacteria with G+C varying from 42% to 67%, and from *Acanthamoeba castellanii*, the tupanvirus host. This was not inhibited by sulbactam or ceftriaxone. RNase activity was estimated to be 0.45±0.15 mU/mg using a fluorescence-based assay. Our results still broaden the range of hosts of MBL fold proteins and demonstrate that such protein can have dual beta-lactamase/nuclease activities. We suggest that they should be annotated according to this finding to avoid further confusion.

## INTRODUCTION

The metallo-beta-lactamase (MBL) superfamily encompasses a large set of enzymes, including MBL and ribonuclease (RNase) Z enzymes^1^. These enzymes are pleitropic proteins that can hydrolyze a wide range of substrates, among which beta-lactams, and DNA or RNA^2,3^. Such capabilities rely on an ancient and highly conserved fold, which represents a stable scaffold that has evolved to perform a broad range of chemical reactions and on which various catalytic, regulatory and structural activities are based^2–4^. This wide array of activities is enabled by variations in the composition and size of loops located near the enzyme active site^3^. A well-known catalytic activity of MBLs consists in breaking beta-lactam rings, which was primarily identified in bacteria^5^. Nevertheless, this hydrolase activity is suspected to have evolved in response to the environmental beta-lactams from an ancestral protein whose function was not related to beta-lactams and which may have been devoid of such hydrolase capability^3^. Concurrently to their capability to interact with various substrates that likely emerged through adaptive evolution, members of the MBLs superfamily have been identified in a broad range of cellular organisms, including bacteria, but also eukaryotes and archaea with a beta-lactamase activity^2,6^.

Giant viruses are *bona fide* microbes as their virions are visible under a light microscope and they display a complexity similar to that of small cellular microorganisms^7,8^. Since their discovery in 2003, their diversity has increased considerably, with nine families and more than 100 isolates cultured. Their classification alongside cellular microorganisms is still debated, but their characteristics clearly distinguish them from conventional viruses^9,10^. We have investigated whether genes encoding members of the MBLs superfamily may also be present in giant viruses. We found one in Tupanvirus deep ocean, a giant mimivirus isolated from Brazilian Atlantic ocean sediments, and confirmed that its product harbored a biologically active MBL fold with both beta-lactamase and RNase activities.

## RESULTS

While annotating the genome of Tupanvirus deep ocean, the second isolate of a new mimivirus genus, *Tupanvirus*^11^, a gene (GenBank: AUL78925.1) that encodes a metallo-hydrolase-like MBL fold was identified (Conserved Protein Domain Family Accession no. cl23716)^12^. This gene has a homolog in the other tupanvirus isolate (Soda Lake) (AUL77644.1). Beyond, best BLASTp hits against cellular organisms included MBL fold harboring proteins from an unclassified deltaproteobacterium whose genome was assembled from a marine water metagenome (evalue, 5e-38; identity, 33.0; coverage, 83%), from an actinobacteria (*Nonnomuraea* spp.) (1e-36; 30.0; 86%), from *Microscilla marina* (6e-34; 28.5%; 89%) and from *Acanthamoeba castellanii* (4e-33; 29.8%; 81%) (***Fig. 1***, ***Supplementary Fig. S1***). Significant BLASTp hits (evalues ranging from 1e−41 to 8e−6) against viruses were also obtained with genes from putative giant viruses whose genomes was assembled from metagenomes obtained from environmental samples^13–15^ and from Cafeteria roenbergensis virus, a distant Mimivirus relative^16^. The 322 amino acid long tupanvirus protein exhibits the conserved MBL motif “HxHxDH” in amino acid positions 60-65. A search for domains using the NCBI conserved domain search (CD Search) tool^17^ identified a MBL fold belonging to a ribonuclease Z (RNase_Z_T_toga, TIGR02650, interval= amino acids 24-273, E-value= 1.81e−14; RNaseZ_ZiPD-like_MBL-fold, cd07717, interval= amino acids 56-282, E-value= 1.63e−04), which is a transfer RNA (tRNA)-processing endonuclease. This Tupanvirus deep ocean protein was further analyzed by three-dimensional comparison against the Phyre2 web portal for protein modeling, prediction and analysis^18^. This analysis reported a best match with 100% confidence and 85% coverage (273 amino acid residues) with the crystal structure of a long form ribonuclease Z (RNase Z) from yeast (template c5mtzA) (***Supplementary Fig. S2***). Proteome analysis conducted for Tupanvirus Soda Lake, and Tupanvirus deep ocean, as previously described^12^, did not allow the detection of these proteins with a MBL fold in the virions. In addition, the dramatic RNA shutdown observed during the replication of this giant virus hindered the achievement of transcriptomic analyses. Interestingly, the genomes of 20 of the 21 (95%) giant viruses found to encode a MBL fold protein concurrently encode tRNAs, whereas this is only the case for 46 of the 122 (38%) giant viruses devoid of a MBL fold protein (p<10^−3^; Yates-corrected chi-square test) (***Supplementary Fig. S3 and 4 and Table S2***). The presence of a MBL fold protein among Megavirales members was correlated with the size of the gene repertoire and the number of translation-associated components (p<10^−3^; Anova test). Putative proteins with a MBL fold from giant viruses comprised two related phylogenetic clusters (***Fig. 1***). These clusters appeared deeply rooted in the phylogenetic tree, which suggests an ancient origin for these genes. In addition, one of the clusters of giant virus genes encoding MBL fold proteins appeared closely related to two genes from *Acanthamoeba castellanii*, an amoebal mimivirus hosts, suggesting a transfer from these giant viruses to *A. castellanii*.

**Figure 1.**
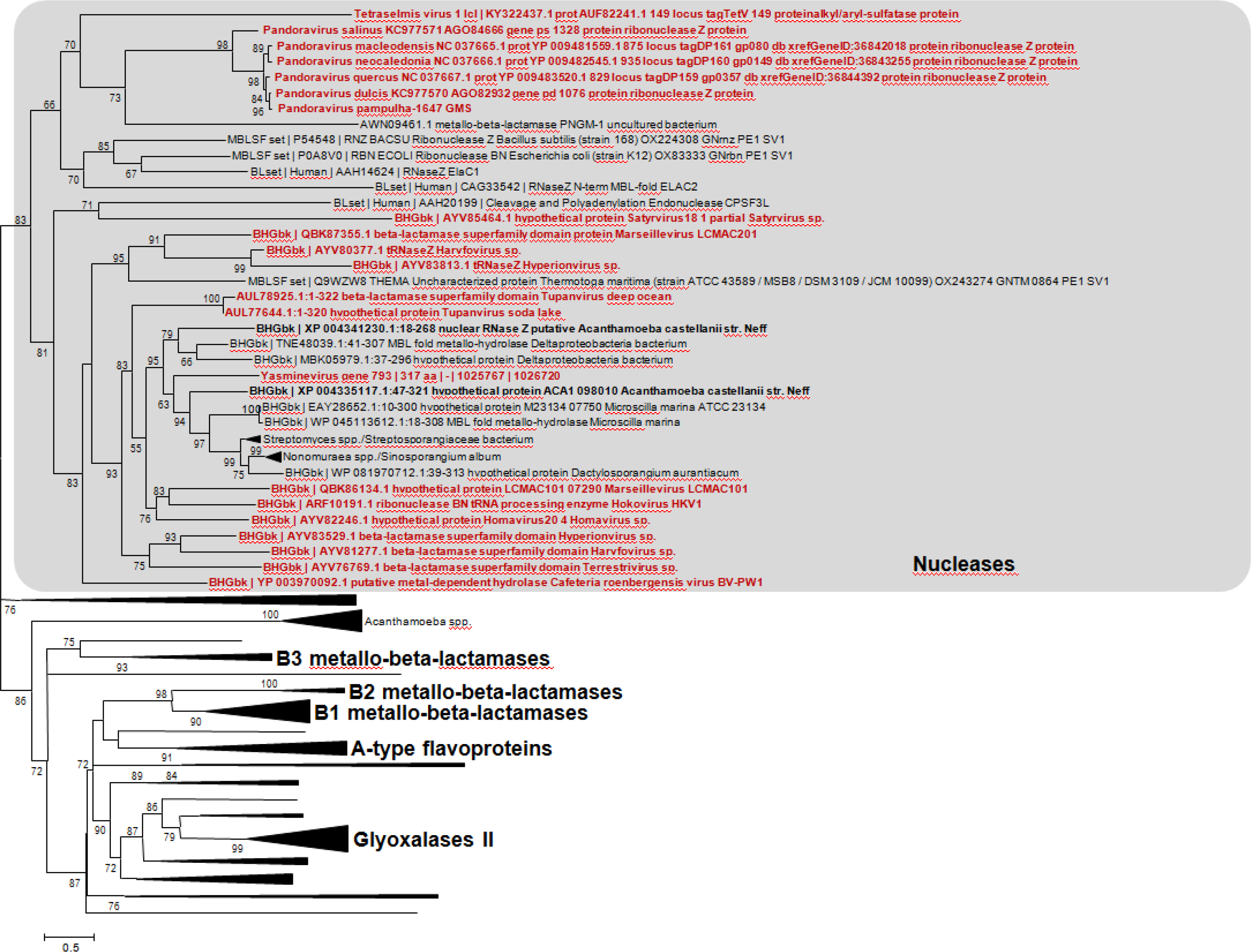
Phylogeny reconstruction based on metallo-beta-lactamase (MBL) fold proteins. Phylogeny reconstruction was performed after amino acid sequence alignment with the Muscle program^33^ with the Maximum-Likelihood method using FastTree^34^, and tree was visualized with the MEGA 6 software^35^. The amino acid sequences analyzed are Tupanvirus deep ocean protein AUL78925.1 and its homologs with the greatest BLASTp scores from the NCBI GenBank protein sequence database (nr) (see ***Supplementary Table S1***), our sequence database of giant virus genomes, and previously described draft genome sequences from 14 *Acanthamoeba* species^36^; a set of previously described MBL fold proteins^19^; and a set of sequences from the UniProtKB database^1^, previously used for phylogeny reconstructions. Extended tree is available in ***Supplementary Figure S1.***

The recombinant protein AUL78925.1 of Tupanvirus deep ocean (named TupBlac) was expressed in *Escherichia coli* and was then purified, as described previously^6^. Based on the phylogenetic analysis and as MBL folds can hydrolyse nucleic acids^2^, both beta-lactamase and nuclease activities of this purified protein were thereafter tested. We first evaluated the beta-lactamase activity of a pure solution of TupBlac used at a concentration of 1 μg/mL by incubating it with nitrocefin, a chromogenic beta-lactam used to test the beta-lactamase activity^19^. A significant hydrolysis activity was observed (***Fig. 2***). A concentrate of protein extract (50 mg/mL) obtained from tupanvirus virions also degraded, albeit slightly, nitrocefin. Thereafter, we monitored by liquid chromatography-mass spectrometry the effect of TupBlac on penicillin G (10 μg/mL) and observed a significant hydrolysis activity of this coumpound within 48h (***Fig. 3***). We also detected, in the presence of the tupanvirus protein, benzylpenilloic acid, the metabolite resulting from the enzymatic hydrolysis of penicillin G^20^. Finally, we confirmed that these observations were related to a beta-lactamase activity as both penicillin G degradation and benzylpenilloic acid appearance were inhibited by sulbactam, a beta-lactamase inhibitor (***Fig. 3***). We further tested if pre-treatment with sulbactam had an impact on the duration of the giant virus replication cycle and replication intensity. After replication on *A. castellanii* strain Neff in the presence of a high concentration (10 μg/mL) of sulbactam, the virions produced (10^6^/mL) were inoculated on fresh amoebae at different concentrations. No differences were observed regarding viral growth in the absence or presence of pre-treatment with sulbactam as assessed using high content screening (***Supplementary Fig. S5***).

**Figure 2:**
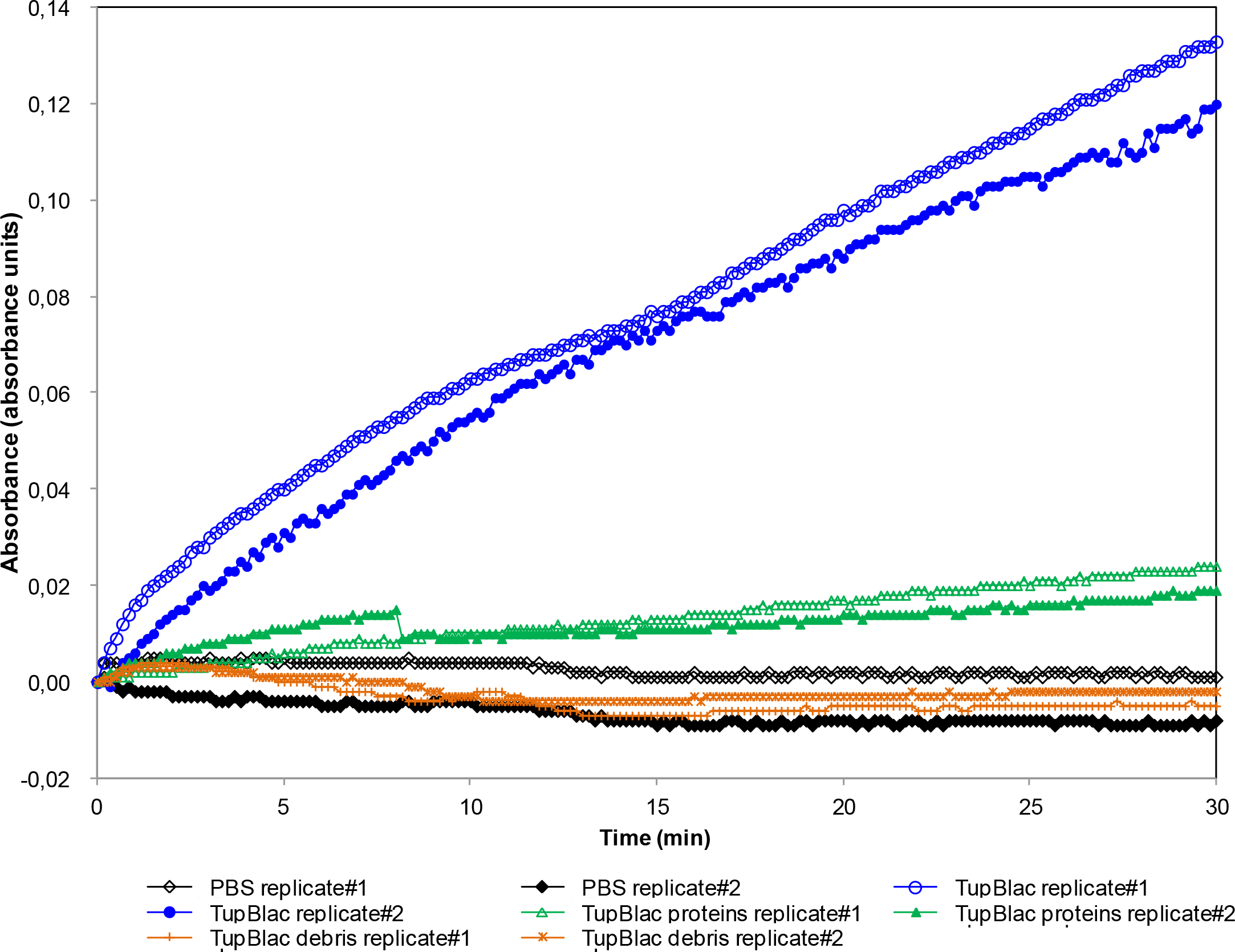
Effect on nitrocefin of expressed Tupanvirus protein (TupBlac). The effect on nitrocefin of the expressed Tupanvirus protein (TupBlac) was assessed by monitoring the degradation of nitrocefin, a chromogenic cephalosporin substrate. PBS, Phosphate-Buffered Saline; TupBlac, tupanvirus expressed protein

**Figure 3:**
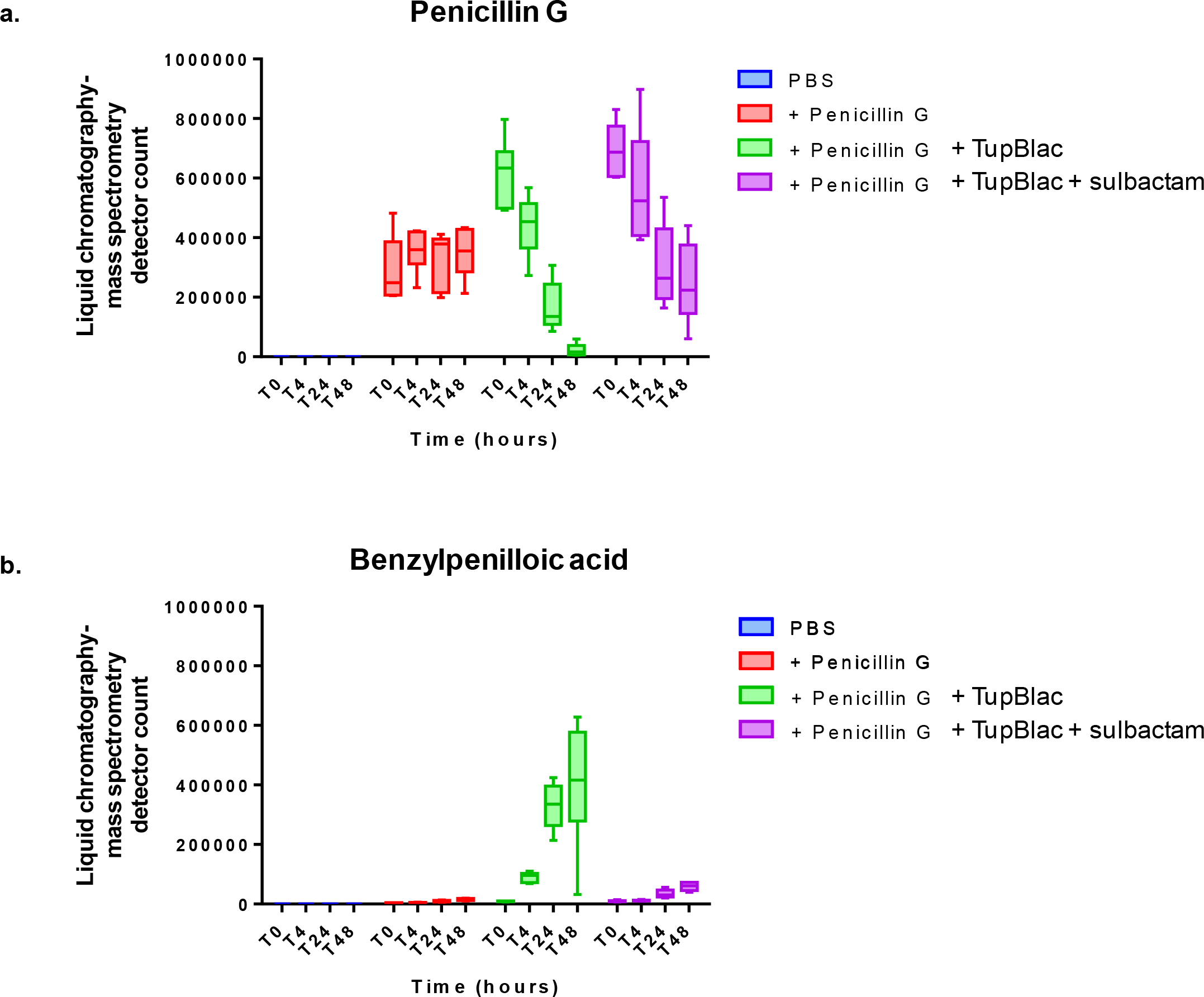
Effect on penicillin G of expressed Tupanvirus protein (TupBlac). The effect on penicillin G of the expressed Tupanvirus protein (TupBlac) and its inhibition by sulbactam were assessed by monitoring by liquid chromatography-mass spectrometry (LC-MS) the degradation of penicillin G (a) and the appearance of benzylpenilloic acid, the metabolite resulting from the enzymatic hydrolysis of penicillin G (b), at times (T) T0 (0 hour), T4 (4 hours), T24 (24 hours), and T48 (48 hours). PBS, Phosphate-Buffered Saline; TupBlac, tupanvirus expressed protein

Finally, as some proteins with a MBL fold can hydrolyse DNA and RNA^2^, we tested the capability of tupanvirus enzyme TupBLac to degrade synthetic single- and double-stranded DNAs and bacterial RNAs. We found no effect on both DNA types. In contrast, we observed a strong RNase activity (***Fig. 4***). Another set of experiments was conducted using *E. coli* RNA as a substrate with an assessment of RNA size distribution on a bioanalyzer (Agilent Technologies, Palo Alto, CA) after incubation with TupBlac. It showed a dramatic degradation of RNAs by the tupanvirus enzyme (***Fig. 5a***). In contrast with the beta-lactamase activity, this was not inhibited, neither by sulbactam (***Fig. 5a*** and ***Supplementary Fig. S6***), nor by ceftriaxone (***Fig. 5b***), a cephalosporin that inhibits human SNM1A and SNM1B, that are DNA repair nucleases with a MBL fold^21^. In addition, a RNase activity of the Tupanvirus protein was further observed on *A. castellanii* RNA, and not inhibited either by sulbactam or ceftriaxone (***Fig. 5c***). TupBLac also degraded RNA extracted from bacteria with genomes with different G+C contents ranging between 41.8% and 66.6% (***Fig. 5d***), suggesting an absence of influence of the G+C richness on the RNase activity. Finally, TupBlac RNase activity was estimated to be 0.451±0.153 mU/mg using a fluorescence-based assay, without difference in the presence of sulbactam or ceftriaxone (0.520±0.003 and 0.551±0.024 mU/mg, respectively) (***Supplementary Fig. S7***).

**Figure 4:**
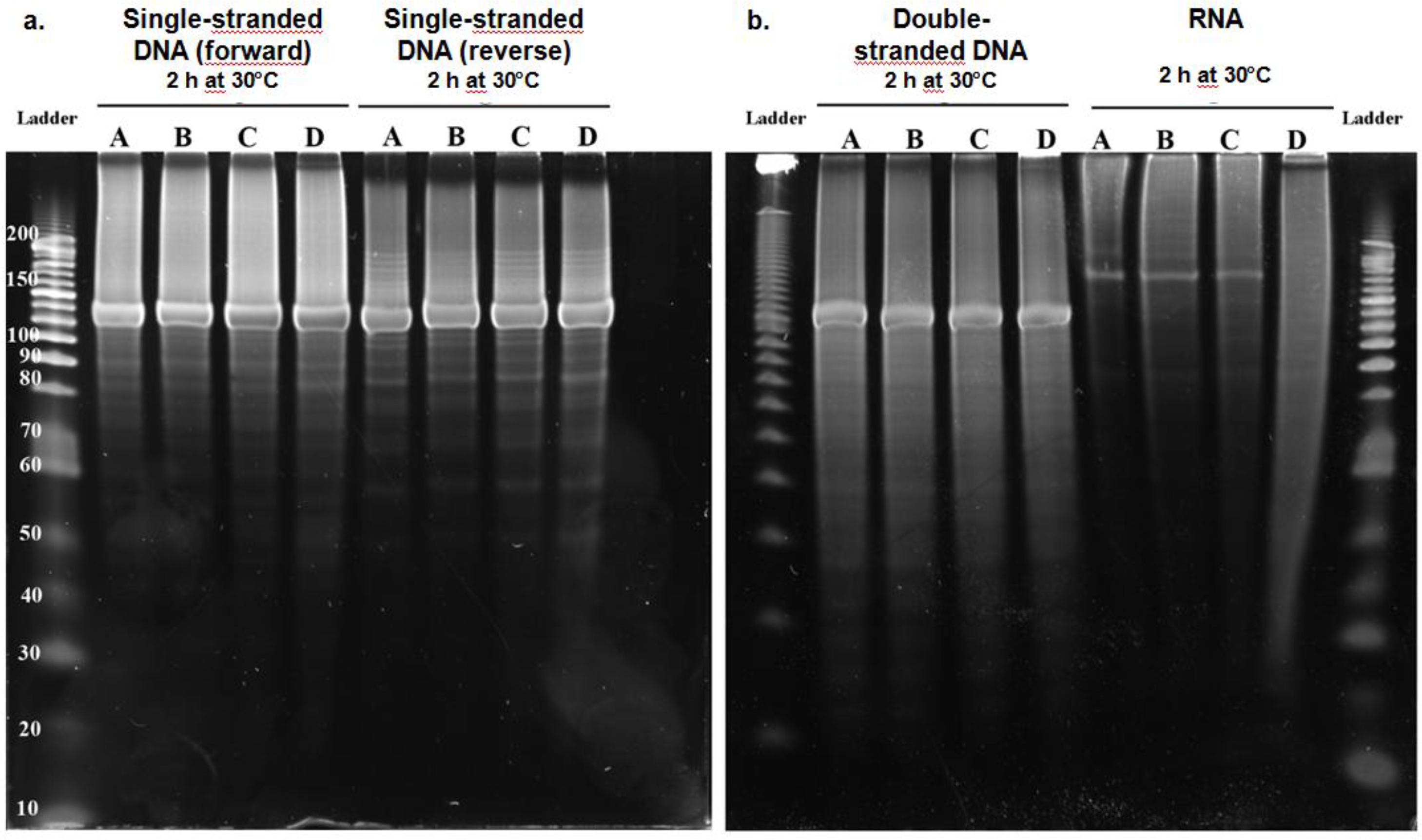
Nuclease activity on various types of nucleic acids of expressed Tupanvirus protein (TupBlac) as assessed by dPAGE. Denaturant polyacrylamide gel electrophoresis (12% dPAGE) of nuclease activity on synthetic (+) and (−) single-stranded DNAs (130 nucleotide-long) (a), synthetic double-stranded DNA (b) (see also ***Supplementary Table S3***),or *Escherichia coli* RNA (b). No treatment (A); buffer (B); succinate dehydrogenase enzyme produced and purified by the same process and collected in the same fractions as Tupanvirus beta-lactamase TupBlac, used as negative control (C); Tupanvirus beta-lactamase TupBlac (D).

**Figure 5:**
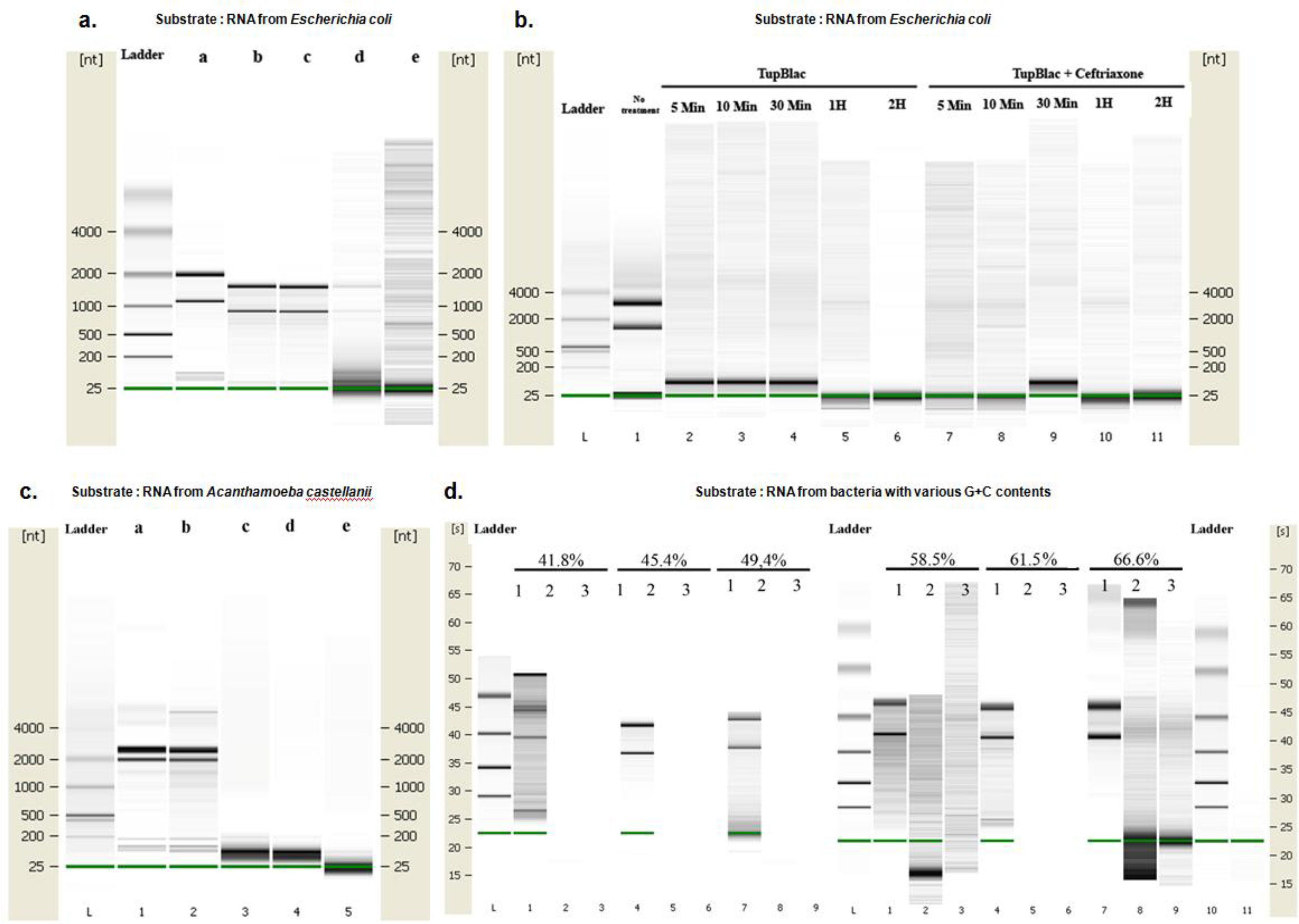
Digital gel images of RNase activity of expressed Tupanvirus protein TupBlac on *E. coli* RNA. RNA samples (1 μg) were incubated with 15 μg of TupBlac at 30°C in the absence or presence of 10 μg/mL of sulbactam or 200 μM of ceftriaxone. Nuclease activity was visualized as digital gel images performed using the Agilent Bioanalyzer 2100 with the RNA 6000 Pico LabChip (Agilent Technologies, Palo Alto, CA). a: RNA used as substrate was from *Escherichia coli;* no treatment (a); buffer (b); sulbactam (c); TupBlac in the absence (d) or presence (e) of sulbactam. b: RNA used as substrate was from *Escherichia coli;* reactions were stopped at different times (5 min, 10 min, 30 min, 1 h and 2 h) by the addition of proteinase K (10 μg) and incubation for 1h at 37°C. The first lane corresponds to no treatment; lanes 2 to 6 to RNA treatment with TupBlac in the absence of ceftriaxone; lanes 7 to 11 to RNA treatment with TupBlac in the presence of ceftriaxone. c: nuclease activity on RNAs originating from *Acanthamoeba castellanii*; no treatment (a); buffer (b); TupBlac in the absence (c, d) or presence (e) of sulbactam. d: nuclease activity on RNAs originating from bacteria that differ by the G+C-content of their genome, as indicated at the top of the digital gel image. For each RNA, three samples were analyzed: no treatment (1); treatment with TupBlac in the absence of sulbactam (2); and treatment with TupBlac in the presence of sulbactam (3).

## DISCUSSION

Hence, we found herein by several bioinformatic approaches that a gene of Tupanvirus deep ocean, a recently discovered giant virus classified in family *Mimiviridae*^11,12^, encodes for a protein with a MBL fold. We further observed that this protein exhibited dual beta-lactamase and RNase activities. This is the first evidence of the presence of a biologically-active protein with a MBL fold in a virus. In addition, this work parallels the one on a protein detected by functional screening of a metagenomic library from the deep-seep sediments^22^, showing that the same enzyme has both beta-lactamase and RNase activities. Indeed, MBL fold proteins were previously biologically-tested for either activity, but not for both. It is noteworthy that the beta-lactamase activity of the MBL fold protein of Tupanvirus was inhibited by a beta-lactamase inhibitor but this was not the case for the RNase activity^23^. The phylogenetic study of this beta-lactamase shows the presence in several other giant viruses of phylogenetically-clustered counterparts, the origin of which seems very old. Interestingly, it also appears that there may have been a gene transfer between these giant viruses and *Acanthamoeba* sp., the amoebal host of many giant viruses. Such potential for horizontal transfer of these MBL fold proteins is well-recognized ^3^.

Beta-lactamases are *a priori* useless for giant viruses, which are grown in the presence of various antibiotics, including beta-lactams^24^, but our findings enhance the recent reconsideration of the function of MBL fold proteins. Thus, the recent description of penicillin secretion by arthropods ^25^ and the demonstration of active beta-lactamase in vertebrates including humans^19^, as well as in archaea^6^ and fungi ^26^ show that MBL fold proteins have a dramatically broad distribution. In humans, 18 genes were annotated as beta-lactamases, whose activity had not been biologically-tested until recently^19^. In addition, MBL fold proteins were highlighted to digest DNA or RNA^2,19^. Thus, a class of enzymes, that were named beta-lactamases because of their original discovery in bacteria resistant to beta-lactamines, are in fact potentially versatile proteins. This differs from the drastically-simplified paradigm consisting in enzymes with a beta-lactamase activity being secreted by bacteria under the selective pressure of natural or prescribed antibiotics.

The RNase activity observed here for the Tupanvirus MBL fold protein could be related to the host ribosomal shutdown observed in the presence of Tupanvirus deep ocean with various protists, the mechanism of which has not been elucidated^12^. This activity could allow these viruses to take over on their cellular hosts by degrading cellular messenger RNAs and shutting down cellular gene expression. The giant virus mRNAs should be protected from such a degradation, which may be explained by the encapsidation of RNA transcripts into giant virions that was detected for some of these viruses^27^. Bioinformatic analyses suggested that the tupanvirus MBL fold protein may belong to the RNase Z group that was proposed to be one of the two main groups of the MBL superfamily with that encompassing MBLs^1^. RNase Z enzymes perform tRNA maturation by catalyzing the endoribonucleolytic removal of the 3’ extension of tRNA precursors that do not contain a chromosomally-encoded CCA determinant^28–30^. The presence in giant viruses of RNases showing the greatest homology to tRNases suggests a specific activity on tRNAs, which seems consistent with the presence of a large set of translation components in these viruses, first and foremost Tupanvirus deep ocean that is the current record holder of the number of translation components (including 70 tRNAs targeting all 20 canonical amino acids). The presence of a putative tRNase in the virus that currently has the most complete set of translation components of the whole virosphere is likely not fortuitous. Furthermore, it was described for *Escherichia coli* that its RNase Z had endoribonucleasic activity on messager RNAs, being responsible for their decay in *in vitro* experiments^29^. This further argues that MBL fold proteins may contain a wide range of activities. PNGM-1, a MBL fold protein whose sequence was recently described from a deep-sea sediment metagenome by detection of its beta-lactamase activity ^31^, was also found to harbor dual beta-lactamase and RNase activities ^22^. MBL fold proteins from giant viruses are clustered with this protein in the phylogenetic analysis. Interestingly, PNGM-1 was suspected to have evolved from a tRNase Z^22^. In conclusion, our data still broaden the range of biological hosts of MBL fold proteins and demonstrate that such proteins can display dual beta-lactamase and nuclease activities. Therefore, we reannotated the tupanvirus MBL fold protein as a beta-lactamase/nuclease.

## MATERIALS AND METHODS

### Bioinformatics

Searches for Tupanvirus deep ocean protein AUL78925.1 homologs were performed using the BLAST tool^32^. Phylogeny reconstruction was performed after amino acid sequence alignment with the Muscle program^33^ and the Maximum-Likelihood method using FastTree^34^, and tree visualization used MEGA 6 software^35^. The amino acid sequences analyzed are Tupanvirus deep ocean protein AUL78925.1 and its homologs with the greatest BLASTp scores from the NCBI GenBank protein sequence database, our sequence database of giant virus genomes, and previously described draft genome sequences from 14 *Acanthamoeba* species^36^; a set of previously described MBL fold proteins^19^; and a set of sequences from the UniProtKB database^1^ previously used for phylogeny reconstructions. Three-dimensional comparisons for protein modeling, prediction and analysis were carried out against the Phyre2 web portal^18^. The set of translation components from each representative of the proposed order Megavirales^37^ was obtained through a BLASTp search^32^ with their repertoire of predicted proteins against clusters of orthologous groups of proteins (COGs) involved in translation (category J)^38^, using 10^−4^ and 50 amino acids as thresholds for e-values and sequence alignment lengths, respectively. The set of tRNAs from each virus was collected using the ARAGORN online tool (http://130.235.244.92/ARAGORN/)^39^. Hierarchical clustering was performed using the MultiExperiment Viewer software^40^ based on the patterns of presence/absence of MBL fold protein, numbers of translation-associated components (number of tRNAs, aminoacyl tRNA-synthetases, other tRNA-associated proteins, other translation-associated proteins) and size of the gene repertoires for Megavirales members (***Supplementary Table S2***). For each item, the maximum value was determined, and values for each virus were considered relatively to these maximum values, being therefore comprised between 0 and 100%.

### Cloning, expression and purification

The Tupanvirus deep ocean gene bioinformatically predicted to encode a beta-lactamase superfamily domain (AUL78925.1^12^) was designed to include a Strep-tag at the N-terminus and optimized for its expression by *Escherichia coli*. It was synthetized by GenScript (Piscataway, NJ, USA) and ligated between the NdeI and NotI restriction sites of a pET24a(+) plasmid. *E. coli* BL21(DE3)-pGro7/GroEL (Takara Shuzo Co., Kyoto, Japan) grown in ZYP-5052 media were used for the expression of the recombinant protein. When the culture reached an O.D._600 nm_= 0.6 at 37°C, the temperature was lowered to 20°C and L-arabinose (0.2% m/v) was added in order to induce the expression of chaperones. After 20 hours, cells were harvested by centrifugation (5,000 g, 30 min, 4°C) and the pellet was resupended in washing buffer (50 mM Tris pH 8, 300 mM NaCl) and then stored at −80°C overnight. Frozen *E. coli* were thawed and incubated on ice for 1 hour after having added lysozyme, DNAse I and PMSF (phenylmethylsulfonyl fluoride) to final concentrations of 0.25 mg/mL, 10 μg/mL and 0.1 mM, respectively. Partially lysed cells were then disrupted by 3 consecutive cycles of sonication (30 seconds, amplitude 45) performed on a Q700 sonicator system (QSonica). Cellular debris were discarded following a centrifugation step (10,000 g, 20 min, 4°C). The Tupanvirus protein was purified with an ÄKTA avant system (GE Healthcare, Bucks, UK) using Strep-tag affinity chromatography (wash buffer: 50 mM Tris pH 8, 300 mM NaCl, and elution buffer: 50 mM Tris pH 8, 300 mM NaCl, 2.5 mM desthiobiotin) on a 5 mL StrepTrap HP column (GE Healthcare). Fractions containing the protein of interest were pooled. Protein purity was assessed using 12.5% SDS-PAGE analysis (Coomassie staining). Protein expression was confirmed by performing MALDI-TOF MS analysis on gel bands previously obtained by SDS-PAGE. Protein concentrations were measured using a Nanodrop 2000c spectrophotometer (Thermo Scientific, Madison, WI, USA).

### Spectrophotometry assay for the detection of beta-lactamase activity in Tupanvirus virions

Tupanvirus purified virions in solution were centrifuged at 5,000 RPM in order to collect 1g of humid matter. Virions were then suspended into 2 mL of a phosphate-buffered saline (PBS) solution at pH 7.4 prepared in water from a commercial salt mixture (bioMerieux, Marcy-l'Etoile, France). Virions were broken after five freeze-thaw cycles followed by 10 minutes of sonication (Q700 sonicator with a Cup Horn, QSonica, Newtown, Connecticut, USA). Integrity of virions was checked by scanning electron microscopy (TM 4000, Hitachi High-Technologie Corporation, Tokyo, Japan). Debris were discarded following a centrifugation step (15,000 g, 10 minutes). The clear supernatant was lyophilized and then reconstituted in 100 μL of PBS (corresponding to a final concentration of 50 mg/mL of soluble proteins). A pure solution of Tupanvirus protein was buffer-exchanged in PBS and the concentration was adjusted to 1 mg/ml. The degradation of nitrocefin (1 mM in PBS), a chromogenic cephalosporin substrate, was monitored as previously described after the addition of virion protein extract or Tupanvirus protein to the solution^6^.

### Beta-lactam antibiotic degradation monitoring by liquid chromatography-mass spectrometry (LC-MS)

Penicillin G and sulbactam stock solutions at 10 mg/mL were freshly prepared in water from the corresponding high purity salts (Sigma Aldrich). A total of 30 μL of tupanvirus protein solution at 1 mg/mL was spiked with penicillin G and sulbactam at a final concentration of 10 μg/mL, before incubation at room temperature. Each time point corresponded to triplicate sample preparations. Negative controls consisted of PBS spiked with penicillin G and sulbactam. Then, 70 μL of acetonitrile were added to each sample, and tubes were vortexed 10 minutes at 16,000 g to precipitate the proteins. The clear supernatant was collected for analysis using an Acquity I-Class UPLC chromatography system connected to a Vion IMS Qtof ion mobility-quadrupole-time of flight mass spectrometer, as previously described ^6^.

### Assessment of the effect of a beta-lactamase inhibitor on Tupanvirus growth

To evaluate the effect of a beta-lactamase inhibitor sulbactam on Tupanvirus growth, we tested Tupanvirus replication on *A. castellanii* pre-incubated with a high dose of sulbactam. Tests were performed in triplicate and amoebae cultivated in trypticase soy medium^12^. Four 1 mL culture wells containing 5.10^5^ *A. castellanii* were incubated at 32°C, one of which contained 500 mg/L of sulbactam. After 24 hours, Tupanvirus was added at a multiplicity of infection (MOI) of 1 in the well with sulbactam. Two other wells were inoculated with Tupanvirus, including one in which 500 mg/L of sulbactam was added. The last well was used as control of amoeba survival. After 24h, amoebae were counted and Tupanvirus was titrated by qPCR as previously described^12^. In order to assess whether sulbactam could have affected newly formed virions, tupanviruses produced on amoebae incubated with sulbactam were inoculated on fresh amoebae at different concentrations. Their growth was monitored using high content screening microscopy every 8h for 48h^41^. Viral replication was compared to that of tupanviruses produced on amoebae non-treated with sulbactam at the same MOIs.

### Nuclease activity assessment

Nuclease activity was assessed using double-stranded DNA, (+) and (−) single-stranded DNAs, and single-stranded RNAs as substrates. Single-stranded DNAs were synthetic polynucleotides (***Supplementary Table S3***); double-stranded DNA was obtained by annealing (+) and (−) single-stranded DNAs in a thermocycler at temperatures decreasing from 95°C to 25°C over 1h. RNAs used as substrate were from *Escherichia coli*, from different bacteria that differ by the G+C content of their genomes (*Streptococcus parasanguinis* (41.8%), *Vibrio parahaemolyticus* (45.4%), *Vitreoscilla massiliensis* (49.4%), *Aeromonas salmonicida* (58.5%), *Aeromonas hydrophila* (61.5%) and *Pseudomonas aeruginosa* (66.6%)), and from *Acanthamoeba castellanii*. RNAs were purified using RNeasy columns (Invitrogen, Carlsbad, CA, USA). Enzymatic reactions were performed by incubating each polynucleotide (2 μg) with 15 μg of the expressed Tupanvirus protein TupBlac in Tris-HCl buffer 50 mM, pH 8.0, sodium chloride 0.3 M, using a final volume of 20 μL at 30°C for 2 h. After incubation, the material was loaded onto denaturing polyacrylamide gel electrophoresis (dPAGE) at 12% or analysed using the Agilent RNA 6000 Pico LabChip kit on an Agilent 2100 Bioanalyzer (Agilent Technology, Palo Alto, CA, USA). Controls were carried out under the same conditions. The action of TupBlac on RNAs was also assayed in the presence of ceftriaxone, an inhibitor of human metallo β-lactamase fold DNA repair nucleases SNM1A and SNM1B^21^. To do this, enzymatic reactions were conducted at 30°C by incubating *E. coli* RNA (1 μg) with TupBlac (15 μg) in the presence of ceftriaxone at 200 μM. At different times, reactions were stopped by addition of proteinase K (10 μg) and incubated 1h at 37°C. For a quantitative assessment of the RNase activity of the TupBlac enzyme, we used the RNaseAlert QC System kit (Fisher Scientific, Illkirch, France) according to the manufacturer's protocol. This assay uses as substrate a fluorescence-quenched oligonucleotide probe that emits a fluorescent signal in the presence of RNase activity. RNase activities were assayed in the absence or presence of sulbactam (10 μg/mL) or ceftriaxone (200 μM). Negative controls were made with all the reagents used (RNase free water, enzyme buffer, sulbactam and ceftriaxone). Fluorescence was monitored continuously at 37°C for 1h by a Synergy HT plate reader (BioTek Instruments SAS, Colmar, France) with a 485/528 nm filter set. RNase activities of TupBlac were estimated using supplied RNase A used as a standard (10 mU/mL). Two independent experiments were conducted.

## Acknowledgments

This work was supported by the French Government under the “Investments for the Future” program managed by the National Agency for Research (ANR), Méditerranée-Infection 10-IAHU-03, and was also supported by Région Provence-Alpes-Côte d’Azur and European funding FEDER PRIMMI (Fonds Européen de Développement Régional - Plateformes de Recherche et d’Innovation Mutualisées Méditerranée Infection). We are thankful to Rania Francis for her technical help.

## Author contributions

Conceived and designed the study: DR, PC, PP, BLS. Designed and/or performed experiments: DR, PC, LP, SA, NA, EC, BLS, PP. Analyzed and interpreted data: DR, PC, EC, BLS, PP. Wrote the manuscript: PC and DR. All authors read and approved the final manuscript.

