## Supplementary information for "A metallo-beta-lactamase with both beta-lactamase and ribonuclease activity is linked with traduction in giant viruses"

**Supplementary Figure S1 | Phylogeny reconstruction based on Tupanvirus deep ocean protein AUL78925.1, its homologs with the greatest BLASTp scores and various MBL superfamily members.**

Phylogeny reconstruction was performed after amino acid sequence alignment with the Muscle program[1] with the Maximum-Likelihood method using FastTree[2], and tree was visualized with the MEGA 6 software[3]. The amino acid sequences analyzed are Tupanvirus deep ocean protein AUL78925.1 and its homologs with the greatest BLASTp scores (see ***Extended Data Table 2***), a set of previously described MBL fold proteins[4] and a set of sequences from the UniProtKB database[5] previously used for phylogeny reconstructions.

**Supplementary Figure S2 | Amino acid alignment and structural prediction obtained using the Phyre2 web portal for protein modeling, prediction and analysis[6] for Tupanvirus deep ocean protein AUL78925.1.**

This figure was copied-pasted from the Phyre2 web portal (http://www.sbg.bio.ic.ac.uk/~phyre2/html/page.cgi?id=index)

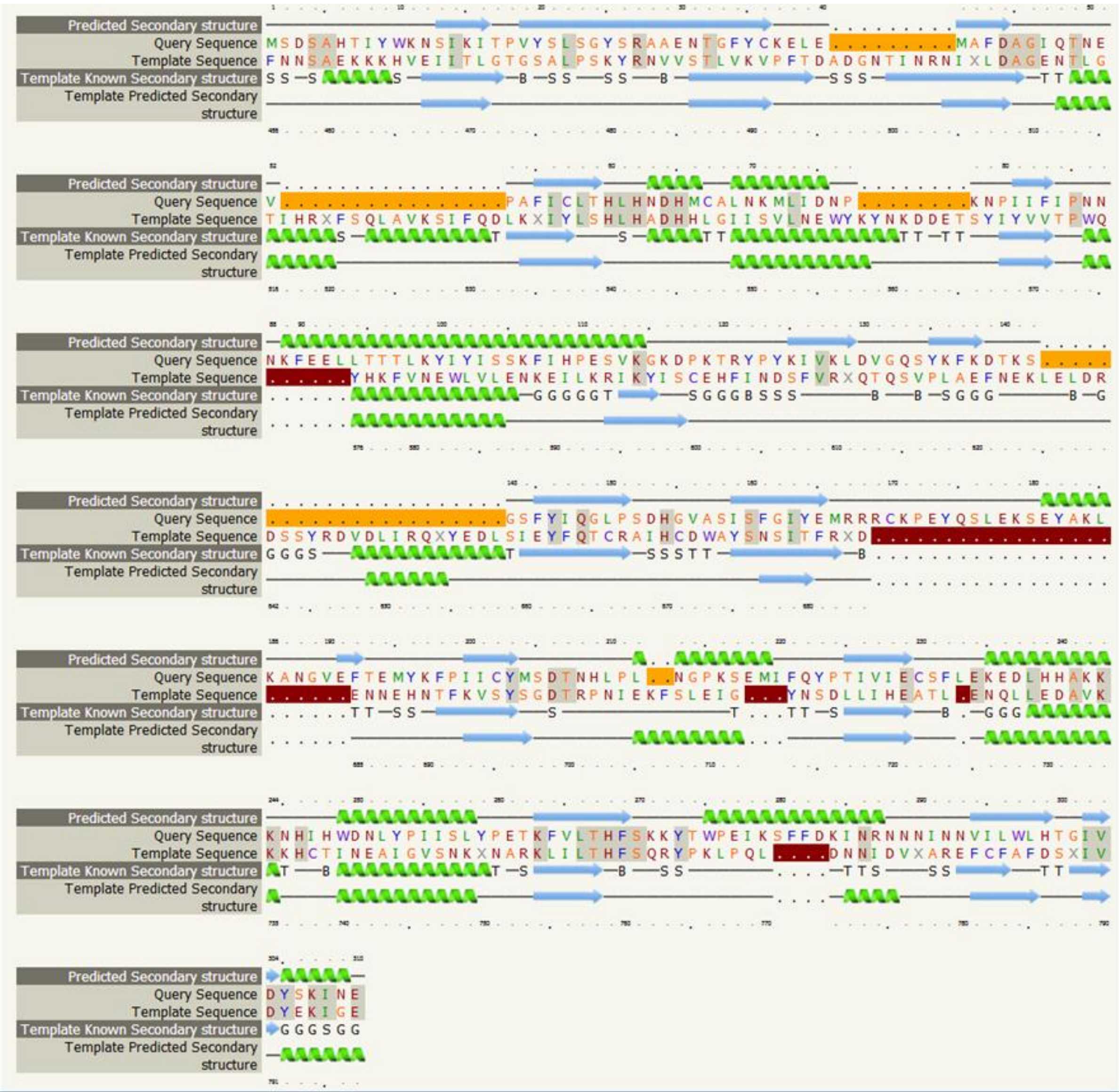

**Supplementary Figure S3 | Number of translation-associated components based on the absence or presence of an MBL fold protein.**

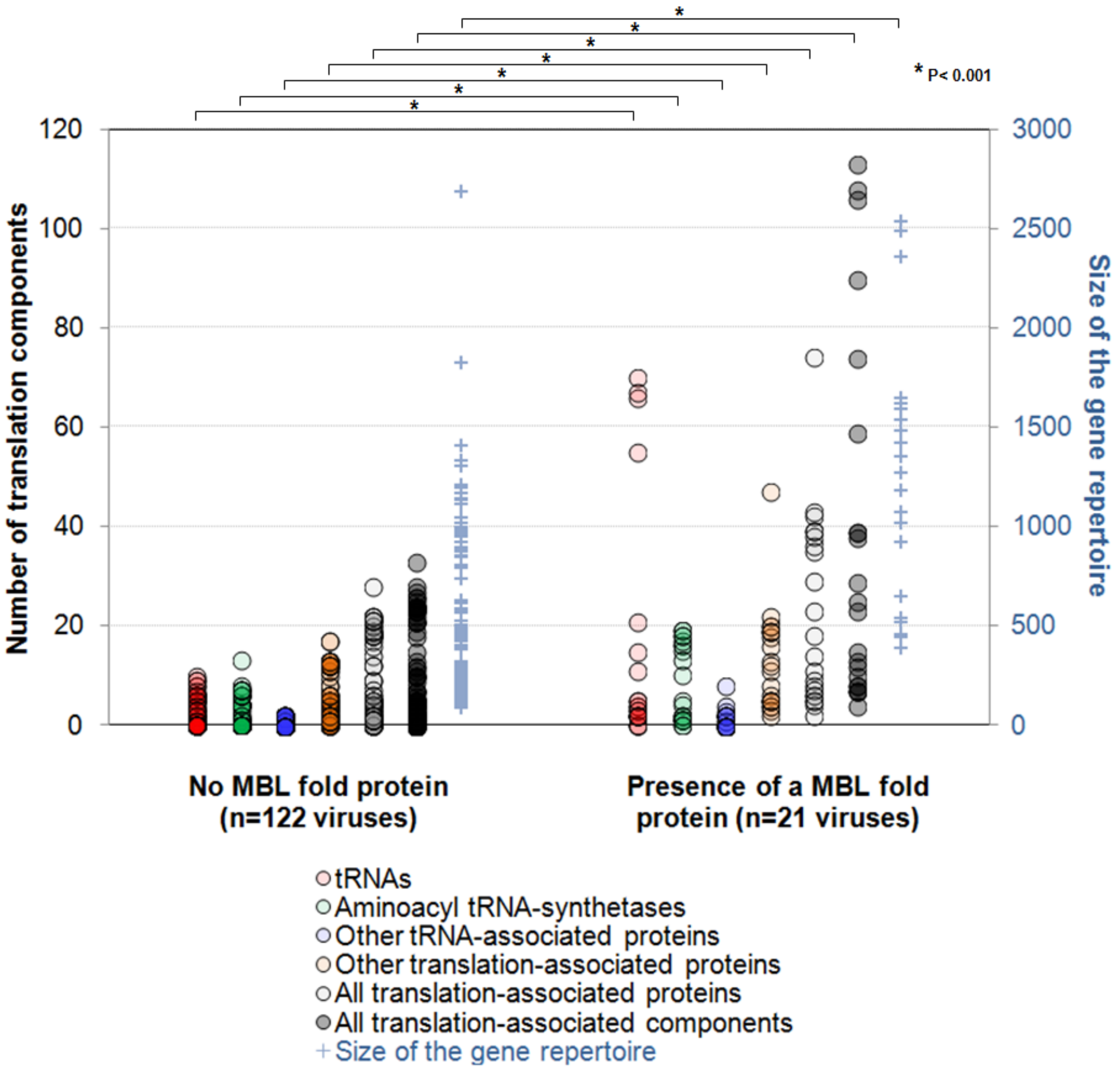

**Supplementary Figure S4 │ Hierarchical clustering based on the presence/absence of an MBL fold protein and other features of giant viruses (see also *Extended Data Table 2*).** Hierarchical clustering was performed using the MultiExperiment Viewer software[7] based on the patterns of presence/absence of MBL fold protein, numbers of translation-associated components (number of tRNAs, aminoacyl tRNA-synthetases, other tRNA-associated proteins, other translation-associated proteins) and size of the gene repertoires for Megavirales members (***Extended Data Table 2***) (a). The MEGA 6 software[3] was used for visualization as a tree (b). For each item, the maximum value was determined, and values for each virus were considered relatively to these maximum values, being therefore comprised between 0 and 100%.

**a.**

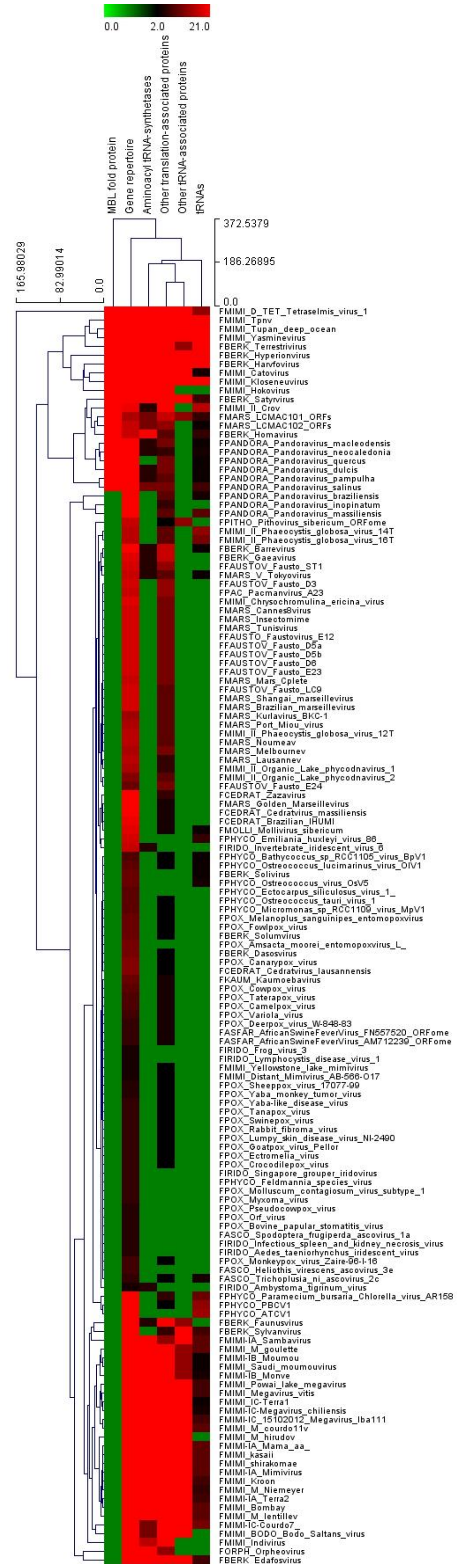
0.0
2.0
21.0
MBL fold protein
Gene repertoire
Aminoacyl tRNA-synthetases
Other translation-associated proteins
Other tRNA-associated proteins
tRNAs
372.5379
186.26895
0.0
165.98029
82.99014
0.0
FMIMI_D_TET_Tetraselmis_virus_1
FMIMI_Tpnv
FMIMI_Tupan_deep_ocean
FMIMI_Yasminevirus
FBERK_Terrestrivirus
FBERK_Hyperionvirus
FBERK_Harvfovirus
FMIMI_Catovirus
FMIMI_Kloseneuvirus
FMIMI_Hokovirus
FBERK_Satyrvirus
FMIMI_II_Crov
FMARS_LCMAC101_ORFs
FMARS_LCMAC102_ORFs
FBERK_Homavirus
FPANDORA_Pandoravirus_macleodensis
FPANDORA_Pandoravirus_neocaledonia
FPANDORA_Pandoravirus_quercus
FPANDORA_Pandoravirus_dulcis
FPANDORA_Pandoravirus_pampulha
FPANDORA_Pandoravirus_salinus
FPANDORA_Pandoravirus_braziliensis
FPANDORA_Pandoravirus_inopinatum
FPANDORA_Pandoravirus_massiliensis
FPITHO_Pithovirus_sibericum_ORFome
FMIMI_II_Phaeocystis_globosa_virus_14T
FMIMI_II_Phaeocystis_globosa_virus_16T
FBERK_Barrevirus
FBERK_Gaeavirus
FFAUSTOV_Fausto_ST1
FMARS_V_Tokyovirus
FFAUSTOV_Fausto_D3
FPAC_Pacmanvirus_A23
FMIMI_Chrysochromulina_ericina_virus
FMARS_Cannes8virus
FMARS_Insectomime
FMARS_Tunisvirus
FFAUSTO_Faustovirus_E12
FFAUSTOV_Fausto_D5a
FFAUSTOV_Fausto_D5b
FFAUSTOV_Fausto_D6
FFAUSTOV_Fausto_E23
FMARS_Mars_Cplete
FFAUSTOV_Fausto_LC9
FMARS_Shangai_marseillevirus
FMARS_Brazilian_marseillevirus
FMARS_Kurlavirus_BKC-1
FMARS_Port_Miou_virus
FMIMI_II_Phaeocystis_globosa_virus_12T
FMARS_Noumeav
FMARS_Melbournev
FMARS_Lausannev
FMIMI_II_Organic_Lake_phycodnavirus_1
FMIMI_II_Organic_Lake_phycodnavirus_2
FFAUSTOV_Fausto_E24
FCEDRAT_Zazavirus
FMARS_Golden_Marseillevirus
FCEDRAT_Cedratvirus_massiliensis
FCEDRAT_Brazilian_IHUMI
FMOLLI_Mollivirus_sibericum
FPHYCO_Emiliania_huxleyi_virus_86_
FIRIDO_Invertebrate_iridescent_virus_6
FPHYCO_Bathycoccus_sp_RCC1105_virus_BpV1
FPHYCO_Ostreococcus_lucimarinus_virus_OIV1
FBERK_Solivirus
FPHYCO_Ostreococcus_virus_OsV5
FPHYCO_Ectocarpus_siliculosus_virus_1_
FPHYCO_Ostreococcus_tauri_virus_1
FPHYCO_Micromonas_sp_RCC1109_virus_MpV1
FPOX_Melanoplus_sanguinipes_entomopoxvirus
FPOX_Fowlpox_virus
FBERK_Solumvirus
FPOX_Amsacta_moorei_entomopoxvirus_L_
FBERK_Dasosvirus
FPOX_Canarypox_virus
FCEDRAT_Cedratvirus_lausannensis
FKAUM_Kaumoebavirus
FPOX_Cowpox_virus
FPOX_Taterapox_virus
FPOX_Camelpox_virus
FPOX_Variola_virus
FPOX_Deerpox_virus_W-848-83
FASFAR_AfricanSwineFeverVirus_FN557520_ORFome
FASFAR_AfricanSwineFeverVirus_AM712239_ORFome
FIRIDO_Frog_virus_3
FIRIDO_Lymphocystis_disease_virus_1
FMIMI_Yellowstone_lake_mimivirus
FMIMI_Distant_Mimivirus_AB-566-O17
FPOX_Sheeppox_virus_17077-99
FPOX_Yaba_monkey_tumor_virus
FPOX_Yaba-like_disease_virus
FPOX_Tanapox_virus
FPOX_Swinepox_virus
FPOX_Rabbit_fibroma_virus
FPOX_Lumpy_skin_disease_virus_NI-2490
FPOX_Goatpox_virus_Pellor
FPOX_Ectromelia_virus
FPOX_Crocodilepox_virus
FIRIDO_Singapore_grouper_iridovirus
FPHYCO_Feldmannia_species_virus
FPOX_Molluscum_contagiosum_virus_subtype_1
FPOX_Myxoma_virus
FPOX_Pseudocowpox_virus
FPOX_Orf_virus
FPOX_Bovine_papular_stomatitis_virus
FASCO_Spodoptera_frugiperda_ascovirus_1a
FIRIDO_Infectious_spleen_and_kidney_necrosis_virus
FIRIDO_Aedes_taeniorhynchus_iridescent_virus
FPOX_Monkeypox_virus_Zaire-96-I-16
FASCO_Heliothis_virescens_ascovirus_3e
FASCO_Trichoplusia_ni_ascovirus_2c
FIRIDO_Ambystoma_tigrinum_virus
FPHYCO_Paramecium_bursaria_Chlorella_virus_AR158
FPHYCO_PBCV1
FPHYCO_ATCV1
FBERK_Faunusvirus
FBERK_Sylvanvirus
FMIMI-IA_Sambavirus
FMIMI_M_goulette
FMIMI-IB_Moumou
FMIMI_Saudi_moumouvirus
FMIMI-IB_Monve
FMIMI_Powai_lake_megavirus
FMIMI_Megavirus_vitis
FMIMI_IC-Terra1
FMIMI-IC-Megavirus_chiliensis
FMIMI-IC_15102012_Megavirus_lba111
FMIMI_M_courdo11v
FMIMI_M_hirudov
FMIMI-IA_Mama_aa_
FMIMI_kasaii
FMIMI_shirakomae
FMIMI-IA_Mimivirus
FMIMI_Kroon
FMIMI_M_Niemeyer
FMIMI-IA_Terra2
FMIMI_Bombay
FMIMI_M_lentillev
FMIMI-IC-Courdo7_
FMIMI_BODO_Bodo_Saltans_virus
FMIMI_Indivirus
FORPH_Orpheovirus
FBERK_Edafosvirus

**b.**

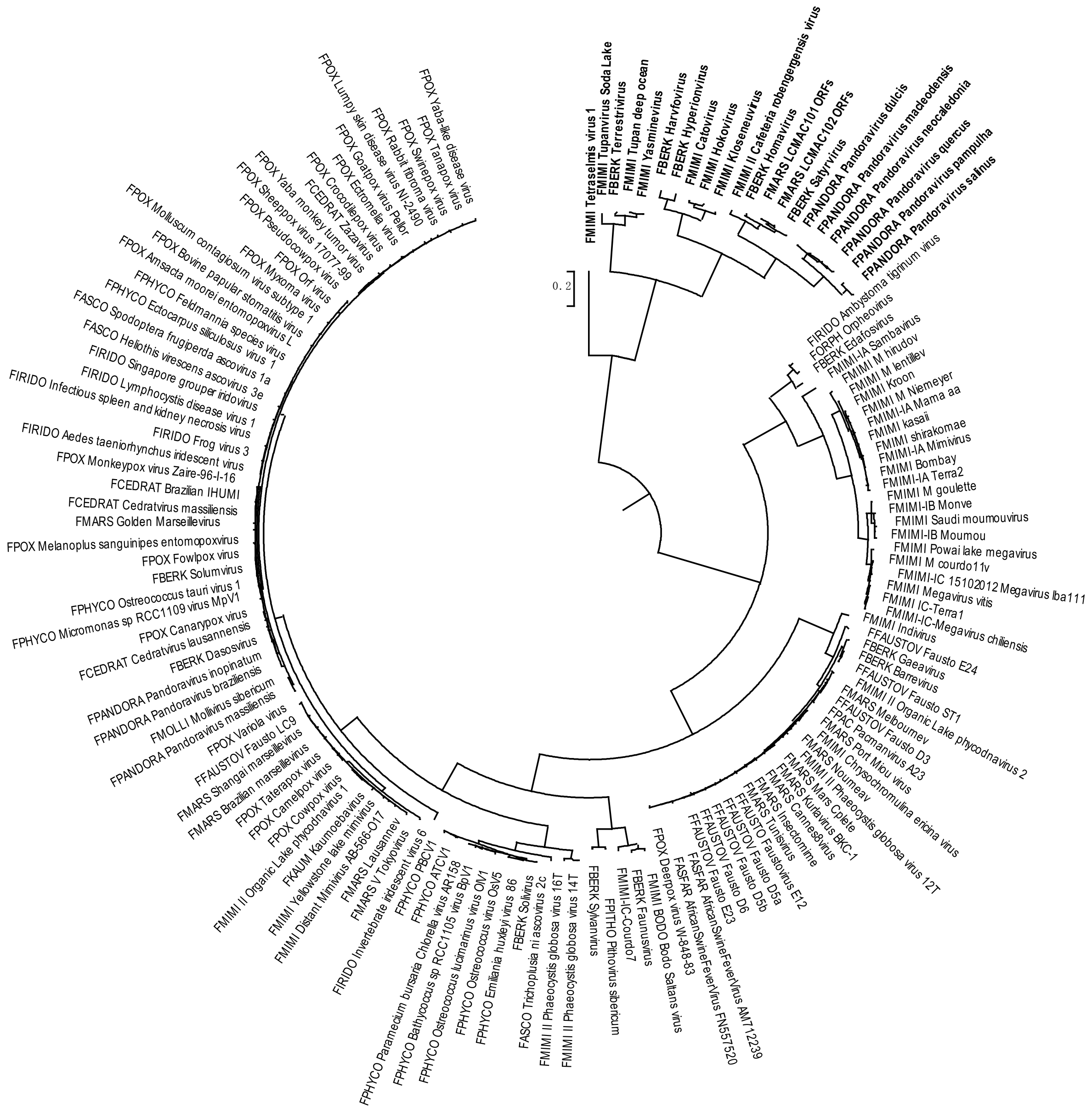
0.2
FMIMI Tetraselmis virus 1
FMIMI Tupanvirus Soda Lake
FBERK Terrestrivirus
FMIMI Tupan deep ocean
FMIMI Yasminevirus
FBERK Harvfovirus
FBERK Hyperionvirus
FMIMI Catovirus
FMIMI Hokovirus
FMIMI Klosneuvirus
FMIMI II Cafeteria roenbergensis virus
FBERK Homavirus
FMARS LCMAC101 ORFs
FMARS LCMAC102 ORFs
FBERK Satyrvirus
FPANDORA Pandoravirus dulcis
FPANDORA Pandoravirus macleodensis
FPANDORA Pandoravirus neocaledonia
FPANDORA Pandoravirus quercus
FPANDORA Pandoravirus pampulha
FPANDORA Pandoravirus salinus
FIRIDO Ambystoma tigrinum virus
FORPH Orpheovirus
FBERK Edafosvirus
FMIMI-IA Sambavirus
FMIMI M hirudov
FMIMI M lentillev
FMIMI Kroon
FMIMI M Niemeyer
FMIMI-IA Mama aa
FMIMI kasaii
FMIMI shirakomae
FMIMI-IA Mimivirus
FMIMI Bombay
FMIMI-IA Terra2
FMIMI M goulette
FMIMI-IB Monve
FMIMI Saudi moumouvirus
FMIMI-IB Moumou
FMIMI Powai lake megavirus
FMIMI M courdo11v
FMIMI-IC 15102012 Megavirus lba111
FMIMI Megavirus vitis
FMIMI IC-Terra1
FMIMI-IC-Megavirus chiliensis
FMIMI Indivirus
FFAUSTOV Fausto E24
FBERK Gaeavirus
FBERK Barrevirus
FFAUSTOV Fausto ST1
FMIMI II Organic Lake phycodnavirus 2
FMARS Melbournev
FFAUSTOV Fausto D3
FPAC Pacmanvirus A23
FMARS Port Miou virus
FMIMI Chrysochromulina ericina virus
FMARS Noumeav
FMIMI II Phaeocystis globosa virus 12T
FMARS Mars Cpiete
FMARS Kurlavirus BKC-1
FMARS Cannes8virus
FMARS Insectomime
FMARS Tunisvirus
FFAUSTOV Faustovirus E12
FFAUSTOV Fausto D5a
FFAUSTOV Fausto D5b
FFAUSTOV Fausto D6
FFAUSTOV Fausto E23
FASFAR AfricanSwineFeverVirus AM712239
FASFAR AfricanSwineFeverVirus FN557520
FPOX Deerpox virus W-848-83
FMIMI BODO Bodo Saltans virus
FBERK Faunusvirus
FMIMI-IC-Courdo7
FPITHO Pithovirus sibericum
FBERK Sylvanvirus
FMIMI II Phaeocystis globosa virus 14T
FMIMI II Phaeocystis globosa virus 16T
FASCO Trichoplusia ni ascovirus 2c
FBERK Solivirus
FPHYCO Emiliania huxleyi virus 86
FPHYCO Ostreococcus virus OsV5
FPHYCO Ostreococcus lucimarinus virus OlV1
FPHYCO Bathycoccus sp RCC1105 virus BpV1
FPHYCO Paramecium bursaria Chlorella virus AR158
FPHYCO ATCV1
FPHYCO PBCV1
FIRIDO Invertebrate iridescent virus 6
FMARS V Tokyovirus
FMARS Lausannev
FMIMI Distant Mimivirus AB-566-O17
FMIMI Yellowstone lake mimivirus
FKAUM Kaumoebavirus
FMIMI II Organic Lake phycodnavirus 1
FPOX Cowpox virus
FPOX Camelpox virus
FPOX Taterapox virus
FMARS Brazilian marseillevirus
FMARS Shangai marseillevirus
FFAUSTOV Fausto LC9
FPOX Variola virus
FPANDORA Pandoravirus massiliensis
FMOLLI Mollivirus sibericum
FPANDORA Pandoravirus braziliensis
FPANDORA Pandoravirus inopinatum
FBERK Dasosvirus
FCEDRAT Cedratvirus lausannensis
FPOX Canarypox virus
FPHYCO Micromonas sp RCC1109 virus MpV1
FPHYCO Ostreococcus tauri virus 1
FBERK Solumvirus
FPOX Fowlpox virus
FPOX Melanoplus sanguinipes entomopoxvirus
FMARS Golden Marseillevirus
FCEDRAT Cedratvirus massiliensis
FCEDRAT Brazilian IHUMI
FPOX Monkeypox virus Zaire-96-I-16
FIRIDO Aedes taeniorhynchus iridescent virus
FIRIDO Frog virus 3
FIRIDO Infectious spleen and kidney necrosis virus
FIRIDO Lymphocystis disease virus 1
FIRIDO Singapore grouper iridovirus
FASCO Heliothis virescens ascovirus 3e
FASCO Spodoptera frugiperda ascovirus 1a
FPHYCO Ectocarpus siliculosus virus 1
FPHYCO Feldmannia species virus
FPOX Amsacta moorei entomopoxvirus L
FPOX Bovine papular stomatitis virus
FPOX Molluscum contagiosum virus subtype 1
FPOX Myxoma virus
FPOX Orf virus
FPOX Pseudocowpox virus
FPOX Sheeppox virus 17077-99
FPOX Yaba monkey tumor virus
FCEDRAT Zazavirus
FPOX Crocodilepox virus
FPOX Ectromelia virus
FPOX Goatpox virus Pellor
FPOX Rabbit fibroma virus
FPOX Lumpy skin disease virus NI-2490
FPOX Swinepox virus
FPOX Tanapox virus
FPOX Yaba-like disease virus

**Supplementary Figure S5 | Tupanvirus deep ocean growth on *Acanthamoeba castellanii* strain Neff after a first passage on *A. castellanii* in the absence (a) or presence (b) of sulbactam.**

Viral growth was assessed by high content screening analysis as described in [8].

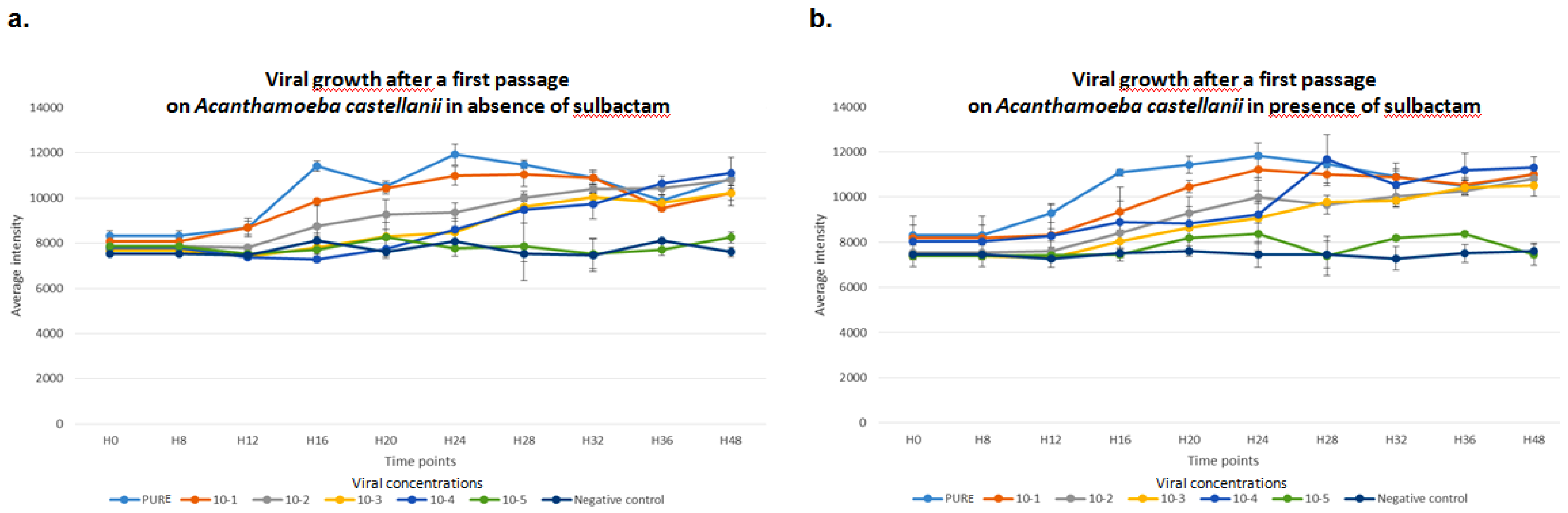

**Supplementary Figure S6 │ RNase activity of expressed Tupanvirus protein TupBlac on *E. coli* RNA, as visualized by a bioanalyzer.**

RNA samples (1 µg) incubated with 15 µg of TupBlac at 30°C in the absence or presence of 10 µg/mL of sulbactam or 200 µM of ceftriaxone. Nuclease activity was visualized as electrophoregrams performed using the Agilent Bioanalyzer 2100 with the RNA 6000 Pico LabChip (Agilent Technologies, Palo Alto, CA). a: no treatment (a); buffer (b); sulbactam (c); TupBlac in the absence (d) or presence (e) of sulbactam. See also Figure 4a for representation from the same data as digital gel image.

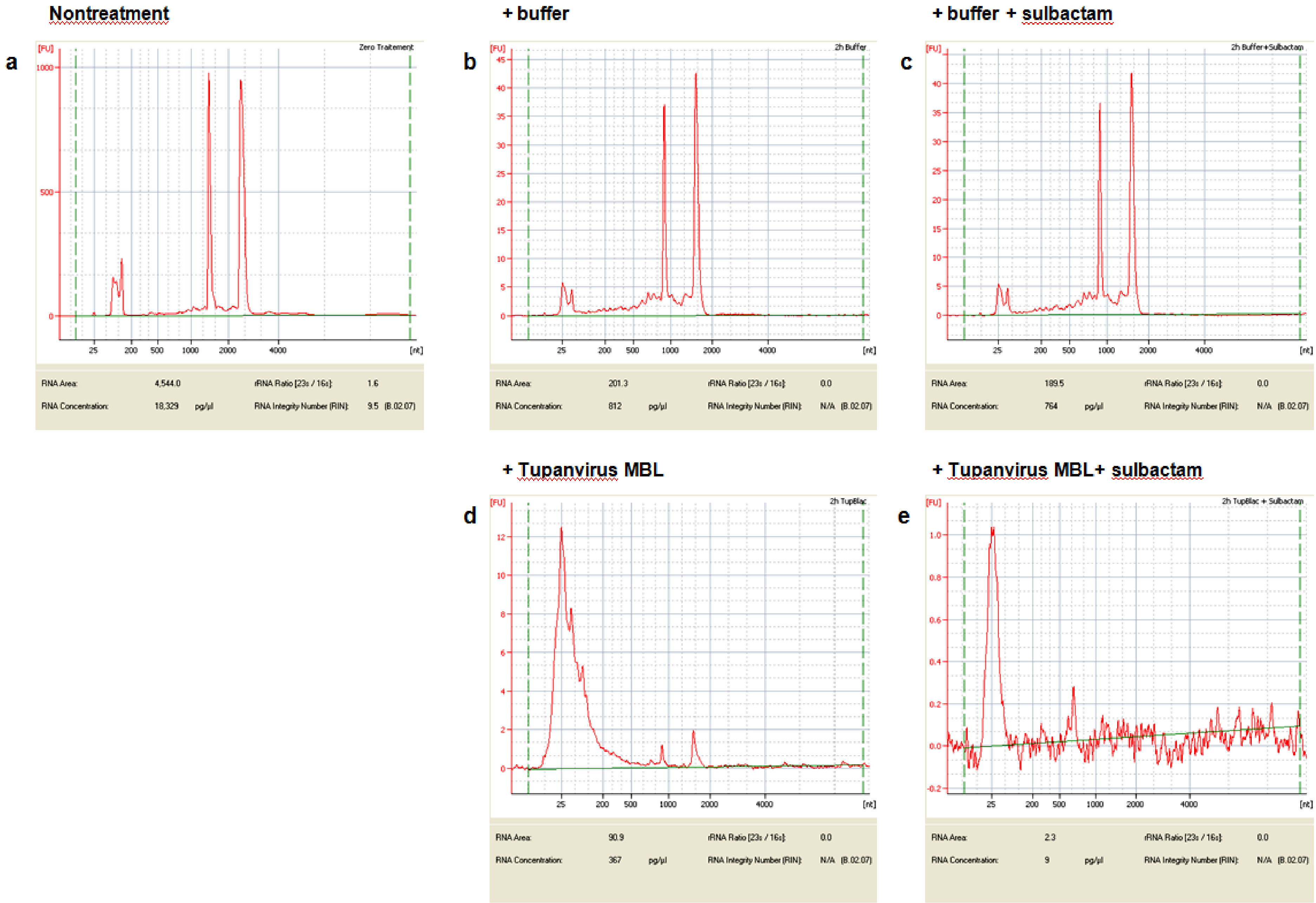
Nontreatment
a
Zero Traitement
RNA Area: 4,544.0
RNA Concentration: 18,329 pg/µl
rRNA Ratio [23s / 16s]: 1.6
RNA Integrity Number (RIN): 9.5 (B.02.07)
+ buffer
b
2h Buffer
RNA Area: 201.3
RNA Concentration: 812 pg/µl
rRNA Ratio [23s / 16s]: 0.0
RNA Integrity Number (RIN): N/A (B.02.07)
+ buffer + sulbactam
c
2h Buffer+Sulbactam
RNA Area: 189.5
RNA Concentration: 764 pg/µl
rRNA Ratio [23s / 16s]: 0.0
RNA Integrity Number (RIN): N/A (B.02.07)
+ Tupanvirus MBL
d
2h TupBlac
RNA Area: 90.9
RNA Concentration: 367 pg/µl
rRNA Ratio [23s / 16s]: 0.0
RNA Integrity Number (RIN): N/A (B.02.07)
+ Tupanvirus MBL+ sulbactam
e
2h TupBlac + Sulbactam
RNA Area: 2.3
RNA Concentration: 9 pg/µl
rRNA Ratio [23s / 16s]: 0.0
RNA Integrity Number (RIN): N/A (B.02.07)
[FU]
[nt]

**Supplementary Figure S7 | Quantitative assessment of RNase activity of expressed Tupanvirus protein TupBlac on *E. coli* RNA.** The RNase activity of TupBlac enzyme was measured using the RNaseAlert™ QC System kit (Fisher Scientific, Illkirch, France) according to the manufacturer's protocol. Fluorescence was monitored continuously at 37°C for 1h in Synergy HT plate reader (BioTek Instruments SAS, Colmar, France) with a 485/528 nm filter set. Two independent experiments were conducted. The addition of TupBlac was associated with a significant increase in fluorescence compared to all controls used (RNase-free water, enzyme buffer, sulbactam, and ceftriaxone). No inhibition of RNase activity of TupBlac was detected with sulbactam or ceftriaxone.

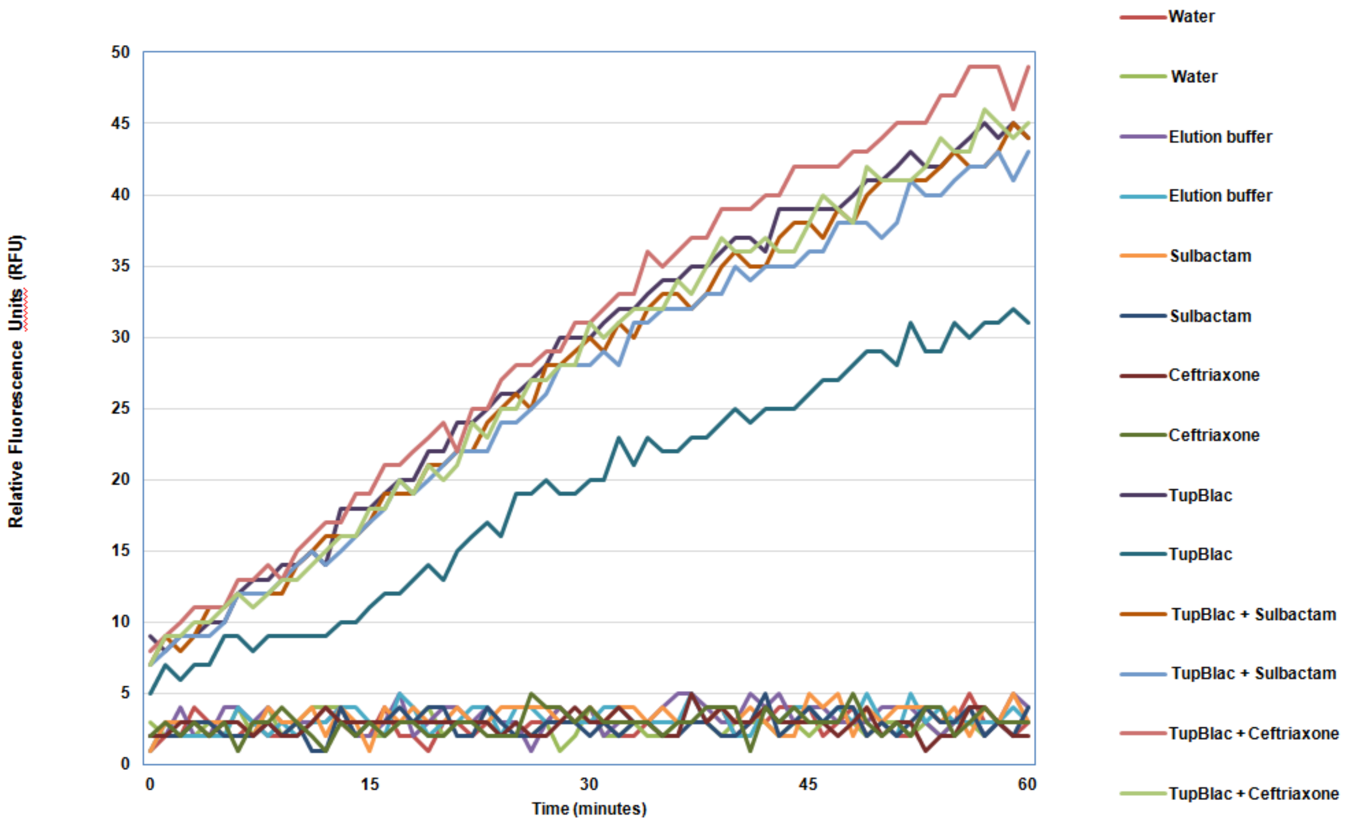

#### Supplementary Table S1 │ Homologs with the 100 greatest BLASTp scores for Tupanvirus deep ocean protein AUL78925.1.

### Query: AUL78925.1 beta-lactamase superfamily domain [Tupanvirus deep ocean]

### 100 hits found

| subject acc.ver | annotation | % identity | alignment length | mismatches | gap opens | q. start | q. end | s. start | s. end | evalue | bit score |
|---|---|---|---|---|---|---|---|---|---|---|---|
| AUL78925.1 | beta-lactamase superfamily domain [Tupanvirus deep ocean] | 100,0 | 322 | 0 | 0 | 1 | 322 | 1 | 322 | 0 | 665 |
| AUL77644.1 | hypothetical protein [Tupanvirus soda lake] | 87,8 | 320 | 39 | 0 | 1 | 320 | 1 | 320 | 0 | 607 |
| TNE48039.1 | MBL fold metallo-hydrolase [Deltaproteobacteria bacterium] | 32,3 | 288 | 165 | 11 | 20 | 298 | 41 | 307 | 1,32E-41 | 156 |
| ARF09172.1 | beta-lactamase superfamily domain [Catovirus CTV1] | 31,0 | 303 | 188 | 8 | 2 | 298 | 7 | 294 | 1,57E-39 | 150 |
| MBK05979.1 | hypothetical protein CL932_144 [Deltaproteobacteria bacterium] | 33,0 | 270 | 170 | 5 | 21 | 289 | 37 | 296 | 5,63E-38 | 147 |
| WP_101786540.1 | MBL fold metallo-hydrolase [Nonomuraea indica] | 30,0 | 287 | 176 | 8 | 21 | 299 | 28 | 297 | 1,59E-36 | 143 |
| AYV76653.1 | beta-lactamase superfamily domain [Terrestrivirus sp.] | 33,2 | 286 | 167 | 8 | 19 | 298 | 36 | 303 | 2,17E-35 | 140 |
| ARF11780.1 | beta-lactamase superfamily domain [Klosneuvirus KNV1] | 31,1 | 283 | 172 | 9 | 20 | 298 | 27 | 290 | 9,67E-35 | 138 |
| WP_090931965.1 | MBL fold metallo-hydrolase [Nonomuraea jiangxiensis] | 27,8 | 299 | 191 | 8 | 10 | 300 | 17 | 298 | 1,11E-34 | 138 |
| EAY28652.1 | hypothetical protein M23134_077 [Microscilla marina ATCC 23134] | 28,5 | 309 | 183 | 8 | 10 | 298 | 10 | 300 | 6,87E-34 | 136 |
| WP_045113612.1 | MBL fold metallo-hydrolase [Microscilla marina] | 28,5 | 309 | 183 | 8 | 10 | 298 | 18 | 308 | 7,94E-34 | 136 |
| WP_132597557.1 | MBL fold metallo-hydrolase [Nonomuraea sp. KC310] | 29,2 | 288 | 179 | 8 | 21 | 300 | 62 | 332 | 8,44E-34 | 137 |
| WP_018384360.1 | hypothetical protein [Streptomyces vitaminophilus] | 26,1 | 303 | 196 | 8 | 10 | 299 | 24 | 311 | 1,05E-33 | 136 |
| XP_004341230.1 | nuclear RNase Z, putative [Acanthamoeba castellanii str. Neff] | 29,8 | 265 | 171 | 7 | 20 | 283 | 18 | 268 | 4,81E-33 | 134 |
| WP_138714153.1 | MBL fold metallo-hydrolase [Nonomuraea sp. 160415] | 28,4 | 285 | 179 | 8 | 21 | 297 | 25 | 292 | 3,13E-32 | 131 |
| WP_106246247.1 | MBL fold metallo-hydrolase [Nonomuraea fuscirosea] | 27,1 | 299 | 194 | 7 | 10 | 300 | 17 | 299 | 3,55E-32 | 131 |
| WP_119925157.1 | hypothetical protein [Streptosporangiaceae bacterium YIM 75507] | 27,4 | 292 | 185 | 9 | 21 | 299 | 35 | 312 | 1,59E-31 | 130 |
| WP_081970712.1 | hypothetical protein [Dactylosporangium aurantiacum] | 28,1 | 292 | 191 | 7 | 10 | 299 | 39 | 313 | 1,59E-31 | 130 |
| WP_138051584.1 | MBL fold metallo-hydrolase [Streptomyces sp. ICN19] | 26,6 | 305 | 194 | 9 | 10 | 299 | 24 | 313 | 2,04E-31 | 130 |
| XP_004335117.1 | hypothetical protein ACA1_0980 [Acanthamoeba castellanii str. Neff] | 29,4 | 289 | 180 | 9 | 20 | 298 | 47 | 321 | 9,72E-31 | 128 |
| WP_014179824.1 | MULTISPECIES: MBL fold metallo-hydrolase [Streptomyces] | 26,4 | 295 | 187 | 9 | 20 | 299 | 34 | 313 | 1,57E-30 | 127 |
| WP_093175703.1 | MBL fold metallo-hydrolase [Sinosporangium album] | 28,2 | 287 | 177 | 8 | 21 | 297 | 25 | 292 | 2,20E-30 | 126 |
| WP_091100978.1 | hypothetical protein [Micromonospora citrea] | 26,5 | 298 | 194 | 8 | 10 | 299 | 40 | 320 | 2,24E-30 | 127 |
| TNE50880.1 | hypothetical protein EP343_069 [Deltaproteobacteria bacterium] | 26,5 | 302 | 197 | 6 | 10 | 299 | 21 | 309 | 3,06E-30 | 127 |
| WP_111832530.1 | MBL fold metallo-hydrolase [Actinomadura madurae] | 26,9 | 305 | 194 | 7 | 10 | 299 | 24 | 314 | 4,83E-30 | 126 |
| WP_075021350.1 | MBL fold metallo-hydrolase [Actinomadura madurae] | 26,9 | 305 | 194 | 7 | 10 | 299 | 24 | 314 | 6,87E-30 | 126 |
| WP_021592586.1 | MBL fold metallo-hydrolase [Actinomadura madurae] | 26,9 | 305 | 194 | 7 | 10 | 299 | 24 | 314 | 7,54E-30 | 125 |
| WP_131737207.1 | MBL fold metallo-hydrolase [Actinomadura roseirufa] | 26,9 | 294 | 186 | 8 | 21 | 299 | 35 | 314 | 1,73E-29 | 125 |
| XP_014160771.1 | hypothetical protein SARC_010 [Sphaeroforma arctica JP610] | 30,7 | 293 | 185 | 8 | 9 | 297 | 10 | 288 | 1,93E-29 | 124 |
| WP_067814206.1 | MBL fold metallo-hydrolase [Actinomadura kijaniata] | 27,2 | 294 | 185 | 9 | 21 | 299 | 34 | 313 | 7,46E-29 | 123 |
| TAE13198.1 | hypothetical protein EAZ95_115 [Bacteroidetes bacterium] | 29,1 | 292 | 180 | 8 | 20 | 300 | 29 | 304 | 9,09E-29 | 122 |
| GAX16660.1 | hypothetical protein FisN_23Lh2 [Fistulifera solaris] | 27,1 | 280 | 193 | 6 | 20 | 296 | 30 | 301 | 9,90E-29 | 123 |
| MBU51739.1 | hypothetical protein CL920_238 [Deltaproteobacteria bacterium] | 26,9 | 294 | 188 | 7 | 20 | 300 | 32 | 311 | 2,13E-28 | 122 |
| OJJ18440.1 | hypothetical protein BKI52_227 [marine bacterium AO1-C] | 25,9 | 294 | 188 | 5 | 21 | 298 | 28 | 307 | 2,45E-28 | 122 |
| WP_109782820.1 | hypothetical protein [Streptomyces sp. CG 926] | 27,5 | 291 | 181 | 9 | 21 | 299 | 35 | 307 | 2,56E-28 | 121 |
| GAX18314.1 | hypothetical protein FisN_23Hh2 [Fistulifera solaris] | 27,1 | 284 | 188 | 6 | 20 | 296 | 55 | 326 | 3,22E-28 | 122 |
| WP_095875954.1 | MBL fold metallo-hydrolase [Streptomyces sp. TLI_235] | 27,1 | 292 | 185 | 10 | 21 | 299 | 35 | 311 | 3,22E-28 | 121 |
| WP_054224620.1 | hypothetical protein [Actinobacteria bacterium OV450] | 26,5 | 302 | 192 | 8 | 10 | 299 | 24 | 307 | 3,65E-28 | 121 |
| WP_089514472.1 | MBL fold metallo-hydrolase [Streptomyces sp. NBS 14/10] | 25,8 | 295 | 189 | 9 | 20 | 299 | 34 | 313 | 5,16E-28 | 121 |
| ARF10191.1 | ribonuclease BN tRNA processing enzyme [Hokovirus HKV1] | 30,5 | 279 | 166 | 4 | 20 | 298 | 18 | 268 | 6,40E-28 | 119 |
| HAS41765.1 | hypothetical protein DCS93_149 [Microscillaceae bacterium] | 26,4 | 296 | 184 | 7 | 21 | 298 | 28 | 307 | 1,33E-27 | 119 |
| PTB75093.1 | hypothetical protein M440DRAFT_13361 [Trichoderma longibrachiatum ATCC 18648] | 27,4 | 296 | 192 | 8 | 22 | 297 | 33 | 325 | 1,47E-27 | 120 |
| RKU37314.1 | metal-dependent hydrolase [Candidatus Poribacteria bacterium] | 31,6 | 269 | 147 | 11 | 21 | 278 | 10 | 252 | 2,32E-27 | 118 |
| KKA29336.1 | hypothetical protein TD95_0019 [Thielaviopsis punctulata] | 25,6 | 317 | 178 | 6 | 20 | 297 | 30 | 327 | 2,36E-27 | 119 |
| WP_052849284.1 | MBL fold metallo-hydrolase [Streptomyces avicenniae] | 26,0 | 304 | 199 | 8 | 10 | 301 | 34 | 323 | 3,43E-27 | 119 |
| WP_030758954.1 | MULTISPECIES: hypothetical protein [unclassified Streptomyces] | 26,5 | 291 | 184 | 9 | 21 | 299 | 35 | 307 | 5,86E-27 | 118 |
| WP_030721416.1 | hypothetical protein [Streptomyces sp. NRRL S-237] | 26,5 | 291 | 184 | 9 | 21 | 299 | 35 | 307 | 6,29E-27 | 118 |
| WP_096330898.1 | MBL fold metallo-hydrolase [Nannocystis exedens] | 25,4 | 295 | 192 | 8 | 20 | 299 | 34 | 315 | 8,68E-27 | 117 |
| XP_009850215.1 | hypothetical protein NEUTE1DRAFT_2430 partial [Neurospora tetrasperma FGSC 2508] | 29,3 | 300 | 181 | 10 | 20 | 298 | 25 | 314 | 9,89E-27 | 117 |
| WP_030277846.1 | MBL fold metallo-hydrolase [Streptomyces sp. NRRL B-24484] | 26,1 | 291 | 188 | 8 | 21 | 299 | 35 | 310 | 1,01E-26 | 117 |

*Supplementary Table S1 - continued*

| # Query: AUL78925.1 beta-lactamase superfamily domain [Tupanvirus deep ocean] | | | | | | | | | | | |
|---|---|---|---|---|---|---|---|---|---|---|---|
| # 100 hits found | | | | | | | | | | | |
| subject acc.ver | annotation | % identity | alignment length | mismatches | gap opens | q. start | q. end | s. start | s. end | evalue | bit score |
| WP_033224288.1 | hypothetical protein [Streptomyces virginiae] | 26,5 | 291 | 184 | 8 | 21 | 299 | 35 | 307 | 1,05E-26 | 117 |
| PNH56094.1 | hypothetical protein VD0003_g15 [Verticillium dahliae] | 27,8 | 299 | 191 | 9 | 20 | 298 | 32 | 325 | 2,08E-26 | 117 |
| WP_150205480.1 | hypothetical protein [Streptomyces venezuelae] | 26,7 | 292 | 183 | 8 | 21 | 299 | 39 | 312 | 2,49E-26 | 116 |
| OAA41047.1 | 3'-tRNA processing endoribonuclease [Metarhizium rileyi RCEF 4871] | 28,6 | 287 | 186 | 7 | 22 | 297 | 32 | 310 | 2,63E-26 | 116 |
| XP_018147478.1 | 3'-tRNA processing endoribonuclease [Pochonia chlamydosporia 170] | 27,2 | 298 | 188 | 8 | 22 | 297 | 32 | 322 | 2,68E-26 | 116 |
| KID90534.1 | RNase (ISS) [Metarhizium guizhouense ARSEF 977] | 28,3 | 297 | 186 | 7 | 22 | 297 | 32 | 322 | 3,22E-26 | 116 |
| WP_026413251.1 | MBL fold metallo-hydrolase [Actinomadura oligospora] | 25,8 | 291 | 188 | 8 | 21 | 299 | 35 | 309 | 3,29E-26 | 116 |
| RZR60587.1 | hypothetical protein I1G_000003 [Pochonia chlamydosporia 123] | 27,5 | 298 | 187 | 8 | 22 | 297 | 32 | 322 | 3,35E-26 | 116 |
| WP_027750155.1 | MBL fold metallo-hydrolase [Streptomyces sp. CNH287] | 25,2 | 306 | 199 | 9 | 10 | 300 | 24 | 314 | 3,64E-26 | 116 |
| WP_030192385.1 | hypothetical protein [Streptomyces sp. NRRL S-87] | 25,8 | 291 | 186 | 8 | 21 | 299 | 37 | 309 | 8,85E-26 | 115 |
| PNH30070.1 | hypothetical protein BJF96_g66 [Verticillium dahliae] | 27,5 | 302 | 188 | 8 | 20 | 298 | 32 | 325 | 9,04E-26 | 115 |
| XP_003348843.1 | uncharacterized protein SMAC_018 [Sordaria macrospora k-hell] | 29,2 | 298 | 185 | 9 | 20 | 298 | 48 | 338 | 9,10E-26 | 115 |
| RKU20277.1 | hypothetical protein C6500_092 [Candidatus Poribacteria bacterium] | 30,6 | 265 | 155 | 8 | 21 | 278 | 10 | 252 | 1,28E-25 | 113 |
| VEU44026.1 | unnamed protein product [Pseudo-nitzschia multistriata] | 27,4 | 270 | 192 | 4 | 21 | 288 | 97 | 364 | 1,45E-25 | 118 |
| PNH48151.1 | hypothetical protein VD0004_g2 [Verticillium dahliae] | 27,5 | 302 | 188 | 8 | 20 | 298 | 32 | 325 | 1,48E-25 | 114 |
| KKF96943.1 | Nuclear ribonuclease [Ceratocystis platani] | 27,0 | 300 | 192 | 5 | 22 | 297 | 32 | 328 | 1,48E-25 | 114 |
| SEG92014.1 | ribonuclease [Nonomuraea solani] | 26,5 | 291 | 183 | 9 | 21 | 299 | 28 | 299 | 2,00E-25 | 114 |
| OLN82381.1 | Nuclear ribonuclease [Colletotrichum chlorophyti] | 29,7 | 300 | 180 | 11 | 20 | 297 | 32 | 322 | 2,14E-25 | 114 |
| XP_956690.1 | hypothetical protein NCU004 [Neurospora crassa OR74A] | 29,0 | 300 | 183 | 10 | 20 | 298 | 49 | 339 | 2,88E-25 | 114 |
| XP_009654568.1 | hypothetical protein VDAG_073 [Verticillium dahliae VdLs.17] | 27,6 | 301 | 187 | 9 | 20 | 297 | 32 | 324 | 2,91E-25 | 114 |
| WP_103958859.1 | hypothetical protein [Nonomuraea solani] | 26,5 | 291 | 183 | 9 | 21 | 299 | 35 | 306 | 3,03E-25 | 113 |
| WP_079403360.1 | hypothetical protein [Streptomyces sp. 3211] | 26,8 | 291 | 183 | 10 | 21 | 299 | 35 | 307 | 3,82E-25 | 113 |
| WP_086748806.1 | MBL fold metallo-hydrolase [Streptomyces scabiei] | 25,9 | 297 | 187 | 7 | 21 | 299 | 50 | 331 | 5,59E-25 | 113 |
| WP_030389651.1 | hypothetical protein [Streptomyces sp. NRRL S-241] | 25,8 | 291 | 186 | 8 | 21 | 299 | 35 | 307 | 5,99E-25 | 112 |
| XP_014549557.1 | RNase (ISS), partial [Metarhizium brunneum ARSEF 3297] | 27,3 | 297 | 189 | 7 | 22 | 297 | 32 | 322 | 8,39E-25 | 112 |
| XP_007808085.1 | 3'-tRNA processing endoribonuclease [Metarhizium acridum CQMa 102] | 27,9 | 294 | 191 | 7 | 22 | 297 | 45 | 335 | 8,81E-25 | 112 |
| WP_060906579.1 | MBL fold metallo-hydrolase [Streptomyces scabiei] | 26,6 | 297 | 185 | 8 | 21 | 299 | 50 | 331 | 1,01E-24 | 112 |
| WP_037702531.1 | MBL fold metallo-hydrolase [Streptomyces scabiei] | 25,9 | 297 | 187 | 7 | 21 | 299 | 50 | 331 | 1,36E-24 | 112 |
| KHN93721.1 | 3'-tRNA processing endoribonuclease [Metarhizium album ARSEF 1941] | 28,5 | 295 | 188 | 8 | 22 | 297 | 32 | 322 | 1,44E-24 | 111 |
| WP_046709922.1 | MBL fold metallo-hydrolase [Streptomyces europaeiscabiei] | 25,9 | 297 | 187 | 7 | 21 | 299 | 50 | 331 | 1,77E-24 | 112 |
| WP_119583429.1 | MBL fold metallo-hydrolase [Streptomyces europaeiscabiei] | 25,9 | 297 | 187 | 7 | 21 | 299 | 50 | 331 | 1,79E-24 | 112 |
| AYV82246.1 | hypothetical protein Homavirus20_4 [Homavirus sp.] | 28,5 | 302 | 187 | 11 | 21 | 308 | 30 | 316 | 1,79E-24 | 111 |
| WP_097249609.1 | MBL fold metallo-hydrolase [Streptomyces sp. 1222.2] | 26,3 | 308 | 194 | 7 | 10 | 299 | 39 | 331 | 1,93E-24 | 111 |
| WP_046914438.1 | MBL fold metallo-hydrolase [Streptomyces stelliscabiei] | 26,3 | 308 | 194 | 7 | 10 | 299 | 39 | 331 | 2,34E-24 | 111 |
| WP_060903867.1 | MBL fold metallo-hydrolase [Streptomyces europaeiscabiei] | 25,9 | 297 | 187 | 7 | 21 | 299 | 50 | 331 | 3,61E-24 | 111 |
| QBK86134.1 | hypothetical protein LCMAC101_072 [Marseillevirus LCMAC101] | 36,7 | 166 | 98 | 5 | 120 | 285 | 42 | 200 | 4,31E-24 | 107 |
| SOD75702.1 | ribonuclease [Streptomyces sp. 1222.2] | 26,3 | 316 | 199 | 8 | 3 | 299 | 76 | 376 | 6,58E-24 | 111 |
| RKU29175.1 | hypothetical protein C6499_089 [Candidatus Poribacteria bacterium] | 29,4 | 265 | 158 | 8 | 21 | 278 | 10 | 252 | 6,92E-24 | 108 |
| ROV91957.1 | hypothetical protein VSDG_076 [Valsa sordida] | 25,9 | 297 | 199 | 6 | 20 | 298 | 30 | 323 | 7,14E-24 | 110 |
| WP_107465387.1 | MBL fold metallo-hydrolase [Streptomyces sp. MA5143a] | 26,4 | 296 | 188 | 7 | 21 | 299 | 35 | 317 | 1,07E-23 | 109 |
| WP_005475892.1 | MBL fold metallo-hydrolase [Streptomyces bottropensis] | 26,3 | 308 | 194 | 7 | 10 | 299 | 39 | 331 | 1,18E-23 | 109 |
| XP_006693126.1 | hypothetical protein CTHT_00266 [Chaetomium thermophilum var. thermophilum DSM 1495] | 27,4 | 314 | 187 | 9 | 21 | 298 | 29 | 337 | 1,27E-23 | 109 |
| WP_055542196.1 | MBL fold metallo-hydrolase [Streptomyces neyagawaensis] | 25,3 | 296 | 191 | 7 | 21 | 299 | 35 | 317 | 1,94E-23 | 108 |
| XP_001908011.1 | uncharacterized protein PODANS_7_65 [Podospora anserina mat+] | 27,6 | 294 | 195 | 9 | 21 | 302 | 47 | 334 | 2,09E-23 | 108 |
| QDU70725.1 | ribonuclease [Planctomycetes bacterium Pan265] | 28,4 | 271 | 147 | 9 | 18 | 272 | 18 | 257 | 2,78E-23 | 107 |
| RKU08627.1 | hypothetical protein C6503_228 [Candidatus Poribacteria bacterium] | 28,7 | 261 | 165 | 7 | 21 | 278 | 10 | 252 | 2,81E-23 | 107 |
| WP_145444879.1 | ribonuclease [Planctomycetes bacterium Pan265] | 28,4 | 271 | 147 | 9 | 18 | 272 | 36 | 275 | 3,48E-23 | 107 |
| VBB86629.1 | Putative protein of unknown function [Podospora comata] | 27,6 | 294 | 195 | 9 | 21 | 302 | 47 | 334 | 3,72E-23 | 108 |
| ORY71307.1 | beta-lactamase-like protein [Pseudomassariella vexata] | 30,4 | 293 | 179 | 8 | 20 | 297 | 32 | 314 | 6,13E-23 | 107 |
| XP_002674945.1 | predicted protein [Naegleria gruberi strain NEG-M] | 28,8 | 274 | 177 | 8 | 18 | 282 | 45 | 309 | 6,45E-23 | 107 |

**Supplementary Table S2 | Presence or absence of a MBL fold protein, size of the gene repertoire, and number of translation-associated components among Megavirales members**

| Virus | Size of the gene repertoire | tRNAs | Aminoacyl tRNA-synthetases | Other tRNA-associated proteins | Other translation-associated proteins | All translation-associated proteins | All translation-associated components |
|---|---|---|---|---|---|---|---|
| **Presence of a MBL fold protein** | | | | | | | |
| FMARS LCMAC101 | 394 | 4 | 2 | 1 | 8 | 11 | 15 |
| FMARS LCMAC102 | 465 | 2 | 2 | 0 | 6 | 8 | 10 |
| FBERK Harvfovirus | 1596 | 15 | 10 | 2 | 11 | 23 | 38 |
| FBERK Hyperionvirus | 2494 | 24 | 16 | 3 | 20 | 39 | 63 |
| FBERK Terrestrivirus | 1652 | 55 | 16 | 1 | 18 | 35 | 90 |
| FMIMI Catovirus | 1427 | 3 | 15 | 2 | 19 | 36 | 39 |
| FMIMI Hokovirus | 1022 | 0 | 13 | 0 | 16 | 29 | 29 |
| FMIMI Kloseneuvirus | 1545 | 21 | 17 | 2 | 19 | 38 | 59 |
| FBERK Homavirus | 451 | 4 | 5 | 0 | 4 | 9 | 13 |
| FMIMI D TET Tetraselmis virus 1 | 653 | 9 | 19 | 8 | 47 | 74 | 83 |
| FMIMI II Crov | 544 | 11 | 1 | 0 | 13 | 14 | 25 |
| FMIMI Tpnv | 1276 | 66 | 18 | 4 | 20 | 42 | 108 |
| FMIMI Tupan deep ocean | 1359 | 70 | 19 | 2 | 22 | 43 | 113 |
| FBERK Satyrvirus | 929 | 5 | 4 | 2 | 12 | 18 | 23 |
| FMIMI YASMV | 1624 | 67 | 18 | 2 | 19 | 39 | 106 |
| FPANDORA Pandoravirus dulcis | 1487 | 2 | 1 | 0 | 5 | 6 | 8 |
| FPANDORA Pandoravirus macleodensis | 926 | 2 | 1 | 0 | 5 | 6 | 8 |
| FPANDORA Pandoravirus neocaledonia | 1081 | 3 | 1 | 0 | 3 | 4 | 7 |
| FPANDORA Pandoravirus pampulha | 2368 | 2 | 1 | 0 | 4 | 5 | 7 |
| FPANDORA Pandoravirus quercus | 1185 | 2 | 0 | 0 | 5 | 5 | 7 |
| FPANDORA Pandoravirus salinus | 2542 | 5 | 2 | 0 | 5 | 7 | 12 |

*Supplementary Table S2 - continued*

| Virus | Size of the gene repertoire | tRNAs | Aminoacyl tRNA-synthetases | Other tRNA-associated proteins | Other translation-associated proteins | All translation-associated proteins | All translation-associated components |
|---|---|---|---|---|---|---|---|
| **No MBL fold protein** | | | | | | | |
| FIRIDO Aedes taeniorhynchus iridescent virus | 126 | 1 | 0 | 0 | 0 | 0 | 1 |
| FIRIDO Ambystoma tigrinum virus | 95 | 0 | 1 | 0 | 0 | 1 | 1 |
| FIRIDO Frog virus 3 | 99 | 0 | 0 | 0 | 0 | 0 | 0 |
| FIRIDO Infectious spleen and kidney necrosis virus | 125 | 0 | 0 | 0 | 0 | 0 | 0 |
| FIRIDO Invertebrate iridescent virus 6 | 468 | 0 | 1 | 0 | 0 | 1 | 1 |
| FIRIDO Lymphocystis disease virus 1 | 110 | 1 | 0 | 0 | 0 | 0 | 1 |
| FIRIDO Singapore grouper iridovirus | 162 | 1 | 0 | 0 | 0 | 0 | 1 |
| FASCO Heliothis virescens ascovirus 3e | 180 | 0 | 0 | 0 | 0 | 0 | 0 |
| FASCO Spodoptera frugiperda ascovirus 1a | 123 | 0 | 0 | 0 | 0 | 0 | 0 |
| FASCO Trichoplusia ni ascovirus 2c | 164 | 3 | 0 | 0 | 1 | 1 | 4 |
| FMARS Brazilian marseillevirus | 488 | 0 | 0 | 0 | 4 | 4 | 4 |
| FMARS Mars Cplete | 457 | 0 | 0 | 0 | 5 | 5 | 5 |
| FMARS Cannes8virus | 483 | 0 | 0 | 0 | 5 | 5 | 5 |
| FMARS Port Miou virus | 410 | 0 | 0 | 0 | 4 | 4 | 4 |
| FMARS Lausannev | 444 | 0 | 0 | 0 | 3 | 3 | 3 |
| FMARS Melbournev | 403 | 0 | 0 | 0 | 5 | 5 | 5 |
| FMARS Insectomime | 477 | 0 | 0 | 0 | 5 | 5 | 5 |
| FMARS Tunisvirus | 484 | 0 | 0 | 0 | 5 | 5 | 5 |
| FMARS Golden Marseillevirus | 592 | 0 | 0 | 0 | 2 | 2 | 2 |
| FMARS Kurlavirus BKC-1 | 386 | 0 | 0 | 0 | 4 | 4 | 4 |
| FMARS Noumeav | 429 | 0 | 0 | 0 | 4 | 4 | 4 |
| FMARS Shangai marseillevirus | 472 | 0 | 0 | 0 | 4 | 4 | 4 |
| FMARS V Tokyovirus | 470 | 2 | 1 | 0 | 4 | 5 | 7 |
| FBERK Gaeavirus | 512 | 0 | 1 | 0 | 8 | 9 | 9 |
| FPITHO Pithovirus sibericum ORFome | 467 | 0 | 0 | 1 | 1 | 2 | 2 |
| FCEDRAT Cedratvirus massiliensis | 574 | 0 | 0 | 0 | 2 | 2 | 2 |
| FCEDRAT Brazilian IHUMI | 533 | 0 | 0 | 0 | 2 | 2 | 2 |
| FCEDRAT Cedratvirus lausannensis | 322 | 0 | 0 | 0 | 1 | 1 | 1 |
| FCEDRAT Zazavirus | 636 | 0 | 0 | 0 | 3 | 3 | 3 |
| FORPH Orpheovirus | 1199 | 1 | 8 | 0 | 6 | 14 | 15 |
| FBERK Solivirus | 282 | 2 | 0 | 0 | 0 | 0 | 2 |
| FBERK Solumvirus | 279 | 0 | 0 | 0 | 1 | 1 | 1 |
| FMIMI BODO Bodo Saltans virus | 1143 | 0 | 2 | 2 | 17 | 21 | 21 |
| FBERK Dasosvirus | 304 | 0 | 0 | 0 | 1 | 1 | 1 |
| FBERK Edafosvirus | 1311 | 5 | 13 | 2 | 13 | 28 | 33 |
| FMIMI Indivirus | 744 | 1 | 3 | 0 | 17 | 20 | 21 |
| FBERK Barrevirus | 587 | 2 | 1 | 0 | 8 | 9 | 11 |
| FMIMI Chrysochromulina ericina virus | 512 | 0 | 0 | 0 | 5 | 5 | 5 |
| FMIMI Distant Mimivirus AB-566-O17 | 120 | 0 | 0 | 0 | 1 | 1 | 1 |
| FMIMI II Organic Lake phycodnavirus 1 | 401 | 0 | 0 | 0 | 3 | 3 | 3 |
| FMIMI II Organic Lake phycodnavirus 2 | 326 | 0 | 0 | 0 | 4 | 4 | 4 |
| FMIMI II Phaeocystis globosa virus 12T | 439 | 0 | 0 | 0 | 4 | 4 | 4 |
| FMIMI II Phaeocystis globosa virus 14T | 433 | 9 | 0 | 0 | 4 | 4 | 13 |
| FMIMI II Phaeocystis globosa virus 16T | 434 | 8 | 0 | 0 | 4 | 4 | 12 |
| FMIMI Yellowstone lake mimivirus | 102 | 0 | 0 | 0 | 1 | 1 | 1 |
| FBERK Faunusvirus | 883 | 0 | 1 | 1 | 10 | 12 | 12 |
| FMIMI kasaii | 988 | 6 | 4 | 2 | 12 | 18 | 24 |
| FMIMI Bombay | 899 | 6 | 4 | 2 | 12 | 18 | 24 |
| FMIMI Kroon | 969 | 5 | 4 | 2 | 12 | 18 | 23 |
| FMIMI shirakomae | 986 | 6 | 4 | 2 | 12 | 18 | 24 |
| FMIMI M goulette | 970 | 6 | 6 | 1 | 11 | 18 | 24 |
| FMIMI M hirudov | 992 | 0 | 4 | 2 | 13 | 19 | 19 |
| FMIMI M lentillev | 807 | 6 | 4 | 2 | 11 | 17 | 23 |
| FMIMI M Niemeyer | 1003 | 4 | 4 | 2 | 12 | 18 | 22 |
| FMIMI-IA Mama aa | 1023 | 6 | 4 | 2 | 13 | 19 | 25 |
| FMIMI-IA Mimivirus | 979 | 6 | 4 | 2 | 12 | 18 | 24 |
| FMIMI-IA Sambavirus | 619 | 6 | 4 | 2 | 6 | 12 | 18 |
| FMIMI-IA Terra2 | 890 | 5 | 4 | 2 | 12 | 18 | 23 |
| FMIMI Saudi moumouvirus | 953 | 2 | 6 | 1 | 12 | 19 | 21 |
| FMIMI-IB Monve | 1150 | 3 | 6 | 1 | 13 | 20 | 23 |
| FMIMI-IB Moumou | 930 | 2 | 6 | 1 | 12 | 19 | 21 |
| FMIMI M courdo11v | 1217 | 6 | 7 | 2 | 13 | 22 | 28 |
| FMIMI Megavirus vitis | 1027 | 5 | 7 | 2 | 12 | 21 | 26 |
| FMIMI Powai lake megavirus | 996 | 5 | 6 | 2 | 13 | 21 | 26 |
| FMIMI IC-Terra1 | 1055 | 2 | 7 | 2 | 13 | 22 | 24 |
| FMIMI-IC 15102012 Megavirus lba111 | 1173 | 5 | 7 | 2 | 13 | 22 | 27 |
| FMIMI-IC-Courdo7 | 1337 | 8 | 2 | 2 | 12 | 16 | 24 |
| FMIMI-IC-Megavirus chiliensis | 1120 | 3 | 7 | 2 | 12 | 21 | 24 |
| FBERK Sylvanvirus | 853 | 5 | 0 | 2 | 3 | 5 | 10 |

*Supplementary Table S2 - continued*

| Virus | Size of the gene repertoire | tRNAs | Aminoacyl tRNA-synthetases | Other tRNA-associated proteins | Other translation-associated proteins | All translation-associated proteins | All translation-associated components |
|---|---|---|---|---|---|---|---|
| **No MBL fold protein** | | | | | | | |
| FPANDORA Pandoravirus braziliensis | 2693 | 3 | 0 | 0 | 4 | 4 | 7 |
| FPANDORA Pandoravirus inopinatum | 1834 | 1 | 0 | 0 | 3 | 3 | 4 |
| FPANDORA Pandoravirus massiliensis | 1414 | 6 | 0 | 0 | 4 | 4 | 10 |
| FMOLLI Mollivirus sibericum | 523 | 2 | 0 | 0 | 2 | 2 | 4 |
| FPHYCO ATCV1 | 860 | 10 | 0 | 0 | 0 | 0 | 10 |
| FPHYCO Bathycoccus sp RCC1105 virus BpV1 | 203 | 2 | 0 | 0 | 1 | 1 | 3 |
| FPHYCO Ectocarpus siliculosus virus 1 | 240 | 0 | 0 | 0 | 0 | 0 | 0 |
| FPHYCO Emiliania huxleyi virus 86 | 472 | 4 | 0 | 0 | 0 | 0 | 4 |
| FPHYCO Feldmannia species virus | 150 | 0 | 0 | 0 | 0 | 0 | 0 |
| FPHYCO Micromonas sp RCC1109 virus MpV1 | 244 | 0 | 0 | 0 | 1 | 1 | 1 |
| FPHYCO Ostreococcus lucimarinus virus OlV1 | 250 | 2 | 0 | 0 | 1 | 1 | 3 |
| FPHYCO Ostreococcus tauri virus 1 | 230 | 0 | 0 | 0 | 1 | 1 | 1 |
| FPHYCO Ostreococcus virus OsV5 | 270 | 3 | 0 | 0 | 0 | 0 | 3 |
| FPHYCO Paramecium bursaria Chlorella virus AR158 | 814 | 7 | 0 | 0 | 3 | 3 | 10 |
| FPHYCO PBCV1 | 802 | 9 | 0 | 0 | 1 | 1 | 10 |
| FFAUSTO Faustovirus E12 | 492 | 0 | 0 | 0 | 5 | 5 | 5 |
| FFAUSTOV Fausto D3 | 485 | 0 | 0 | 0 | 6 | 6 | 6 |
| FFAUSTOV Fausto D5a | 490 | 0 | 0 | 0 | 5 | 5 | 5 |
| FFAUSTOV Fausto D5b | 491 | 0 | 0 | 0 | 5 | 5 | 5 |
| FFAUSTOV Fausto D6 | 495 | 0 | 0 | 0 | 5 | 5 | 5 |
| FFAUSTOV Fausto E23 | 495 | 0 | 0 | 0 | 5 | 5 | 5 |
| FFAUSTOV Fausto E24 | 278 | 0 | 0 | 0 | 5 | 5 | 5 |
| FFAUSTOV Fausto LC9 | 477 | 0 | 0 | 0 | 4 | 4 | 4 |
| FFAUSTOV Fausto ST1 | 478 | 0 | 1 | 0 | 6 | 7 | 7 |
| FKAUM Kaumoebavirus | 264 | 0 | 0 | 0 | 2 | 2 | 2 |
| FPAC Pacmanvirus A23 | 465 | 1 | 0 | 0 | 6 | 6 | 7 |
| FASFAR AfricanSwineFeverVirus AM712239 | 156 | 0 | 0 | 0 | 2 | 2 | 2 |
| FASFAR AfricanSwineFeverVirus FN557520 | 163 | 0 | 0 | 0 | 2 | 2 | 2 |
| FPOX Amsacta moorei entomopoxvirus L | 294 | 0 | 0 | 0 | 0 | 0 | 0 |
| FPOX Bovine papular stomatitis virus | 131 | 0 | 0 | 0 | 0 | 0 | 0 |
| FPOX Camelpox virus | 211 | 0 | 0 | 0 | 2 | 2 | 2 |
| FPOX Canarypox virus | 328 | 0 | 0 | 0 | 1 | 1 | 1 |
| FPOX Cowpox virus | 233 | 0 | 0 | 0 | 2 | 2 | 2 |
| FPOX Crocodilepox virus | 173 | 0 | 0 | 0 | 1 | 1 | 1 |
| FPOX Deerpox virus W-848-83 | 169 | 0 | 0 | 0 | 2 | 2 | 2 |
| FPOX Ectromelia virus | 173 | 0 | 0 | 0 | 1 | 1 | 1 |
| FPOX Fowlpox virus | 261 | 0 | 0 | 0 | 1 | 1 | 1 |
| FPOX Goatpox virus Pellor | 150 | 0 | 0 | 0 | 1 | 1 | 1 |
| FPOX Lumpy skin disease virus NI-2490 | 156 | 0 | 0 | 0 | 1 | 1 | 1 |
| FPOX Melanoplus sanguinipes entomopoxvirus | 267 | 0 | 0 | 0 | 1 | 1 | 1 |
| FPOX Molluscum contagiosum virus subtype 1 | 163 | 0 | 0 | 0 | 0 | 0 | 0 |
| FPOX Monkeypox virus Zaire-96-I-16 | 191 | 0 | 0 | 0 | 1 | 1 | 1 |
| FPOX Myxoma virus | 170 | 0 | 0 | 0 | 0 | 0 | 0 |
| FPOX Orf virus | 130 | 0 | 0 | 0 | 0 | 0 | 0 |
| FPOX Pseudocowpox virus | 134 | 0 | 0 | 0 | 0 | 0 | 0 |
| FPOX Rabbit fibroma virus | 165 | 0 | 0 | 0 | 1 | 1 | 1 |
| FPOX Sheeppox virus 17077-99 | 148 | 0 | 0 | 0 | 1 | 1 | 1 |
| FPOX Swinepox virus | 150 | 0 | 0 | 0 | 1 | 1 | 1 |
| FPOX Tanapox virus | 156 | 0 | 0 | 0 | 1 | 1 | 1 |
| FPOX Taterapox virus | 225 | 0 | 0 | 0 | 2 | 2 | 2 |
| FPOX Variola virus | 197 | 0 | 0 | 0 | 2 | 2 | 2 |
| FPOX Yaba monkey tumor virus | 140 | 0 | 0 | 0 | 1 | 1 | 1 |
| FPOX Yaba-like disease virus | 152 | 0 | 0 | 0 | 1 | 1 | 1 |

**Supplementary Table S3 | Sequence of synthetic DNA (130 nucleotides) used in enzyme treatments as substrates as a single strand or double strand obtained by annealing forward with reverse DNA synthetic.**

| Name | Sequence |
|---|---|
| DNA_Synt_Fwd | ATAGAACAACCAAAAAAAATATCAAAATCTAGAGATGAATCAAGTGAATCA<br>GAAGAATCTGATAATGAATCTGATAATGAATCCGATGAGGAAGTTGAATCA<br>GAAACTGAGATAGAACCAGTCAAATCTAA |
| DNA_Synt_Rev | TTAGATTTGACTGGTTCTATCTCAGTTTCTGATTCAACTTCCTCATCGGATTC<br>ATTATCAGATTCATTATCAGATTCTTCTGATTCACTTGATTCATCTCTAGATT<br>TTGATATTTTTTTGGTTGTTCTAT |
